# Toroidal topology of grid-cell activity precedes spatial navigation during development

**DOI:** 10.64898/2026.03.10.710908

**Authors:** Matteo Guardamagna, Erik Hermansen, Jordan Carpenter, Christine M. Lykken, Benjamin A. Dunn, Edvard I. Moser, May-Britt Moser

## Abstract

The medial entorhinal cortex (MEC) is a central component of the mammalian navigation system^1–5^, in which spatially and directionally tuned neurons, including grid cells and head-direction cells, encode an animal’s position and orientation^6–8^. These cells form internal maps with periodic topologies: head-direction cells traverse ring-like manifolds^9,10^, while grid cells are organized on toroidal manifolds^11^. The persistence of these topologies across behavioral states and environments^11,12^ raises the possibility that they arise intrinsically from network architecture rather than through sensory experience^4,5,13–17^. Consistent with this view, observations in juvenile rats have shown that rudimentary spatial tuning appears in place cells, head direction cells and grid cells almost as soon as pups begin to explore their surroundings at 2-3 weeks of age^18–20^. However, it remains unclear whether spatial experience is required for the initial emergence of positional tuning and for the organization of tuned cells into periodic maps. Here we show, using large-scale ensemble recordings in rat pups, that toroidal manifolds emerge in MEC subnetworks as early as postnatal day 10 (P10), preceding eye and ear opening, upright posture, quadrupedal gait, and active exploration^21,22^. These toroidal networks were modular from the beginning, with increasing differentiation of their dynamics appearing on P11–12. The onset of toroidal topology coincided with a transition in MEC network activity characterized by desynchronization and increased inhibitory connectivity, a developmental shift observed broadly across cortical regions at this age^23,24^. In contrast, ring-like manifolds were already detectable by P9, with traces of directional tuning appearing in individual cells at P8 — consistent with an earlier maturation of subcortical circuitry^25^. As pups subsequently began to explore the environment around P15–16, these internally generated maps progressively aligned with external landmarks, culminating in stable, periodic firing fields by three weeks of age. Taken together, these findings identify ring-like and toroidal manifolds as instinctive computational motifs of the developing brain. Their early emergence, preceding major sensory input and navigation, supports the view that spatial representations are preconfigured and later anchored to the external world through experience-dependent plasticity.

## Main Text

The ability to navigate through space is fundamental to survival and therefore expected to emerge early in life. In adult mammals, spatial navigation relies on circuits centered in the hippocampal and parahippocampal cortices, which together support a cognitive map of the environment^4,26^. Within these circuits, distinct neural populations encode complementary aspects of space: *place cells* fire at specific locations^27^, *head direction cells*^28^ and *internal direction cells*^29^ signal the animaĺs orientation, and *grid cells* exhibit a periodic, lattice-like firing pattern that tiles the environment^6,7^.

In developing rodents, these spatially tuned cells adopt adult-like firing patterns soon after the onset of spatial exploration. Semi-stable tuning in head direction cells, place cells, and border cells emerges by P15–16, whereas grid cells undergo a more protracted maturation, acquiring adult-like hexagonal organization in space between P20 and P30^18–20,30–32^. Whether this developmental progression reflects the gradual, experience-dependent construction of individual spatial maps or alternatively the anchoring of preconfigured internal networks with external space remains unresolved. The observation that the coactivity patterns among head direction cell pairs are already stable before eye opening^20,33,34^, around P12–14, raises the possibility that key elements of spatial network architecture are established before quadrupedal gait and spatial experience. Yet, at such early developmental stages, assessing the presence of internal spatial maps, in the absence of locomotion, requires access to the collective activity of large neural populations^35^, a challenge that has only recently become tractable with the advent of high-density recording technologies.

The present study was designed to determine whether spatial maps are present prior to spatial experience. We used large-scale Neuropixels recordings^36^ to examine neural population activity in the MEC and parasubiculum (PaS) of rat pups before the onset of exploratory locomotion. Most recordings were performed between postnatal days P8 to P12, when pups lack the postural control required for coordinated upright gait, their eyes and ear canals remain closed, and they spend most of their time asleep in the nest (Fig. 1a)^21,37^. Additional recordings were obtained from P15 onward, after the onset of spatial exploration. Because the youngest pups we tested do not yet move extensively through space, their neural activity cannot be meaningfully linked to physical position, preventing the use of conventional spatial analyses to check the expression of grid and other functional cells. However, this constraint provides a unique opportunity to determine whether internal spatial organization emerges intrinsically, in the absence of external spatial input. By inferring map structure from the coactivity patterns of entorhinal neural ensembles, we show that the ring-like topology of head direction representations^9^ and the toroidal topology of grid-cell representations^11^ are expressed before spatial experience, and only later, through navigational behavior, become anchored to the external environment.

**Figure 1.**
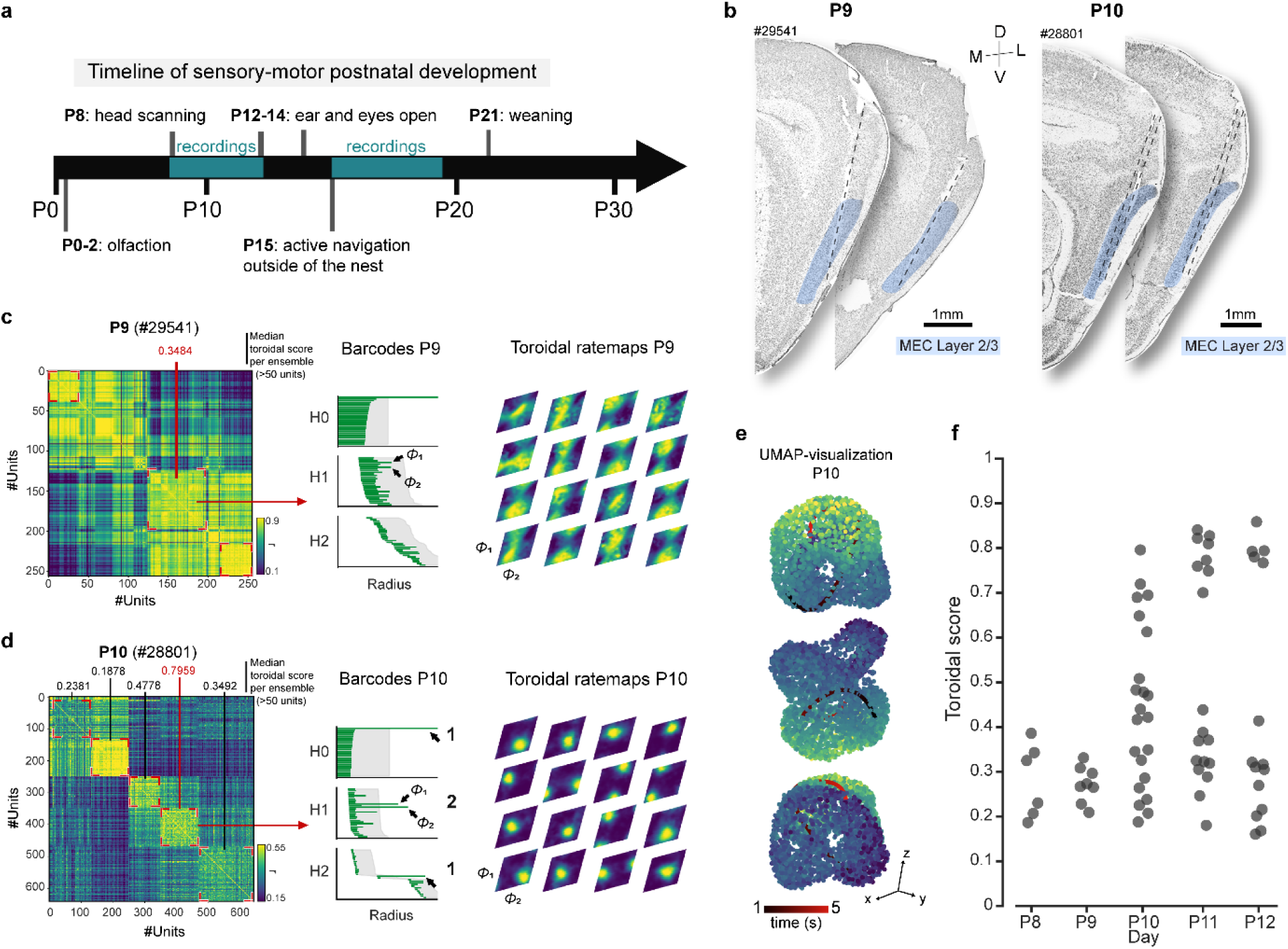
Toroidal manifolds precede onset of navigation. a) Schematic timeline of sensory-motor postnatal development in the rat. Approximate ages for the emergence of key behavioral and sensory-motor functions are indicated. b) Nissl-stained sagittal sections from two rat pups, P9 and P10 (#29541 and #28801, respectively), implanted in MEC Layer 2/3 with Neuropixels 2.0 probe and with tracks highlighted by dotted lines. Sections are shown from lateral (left) to medial (right). MEC Layer 2/3 is highlighted in light blue. c) Time-lagged crosscorrelogram showing correlations between all pairs of MEC-PaS neurons recorded simultaneously in one session at P9 (left). For each cell pair, the ratio between minimum and maximum correlation at time lags from 0 to 300 ms (bins of 10 ms) was used to define the association of the cell pairs. Cell pairs are grouped into clusters of co-active cells, visible as square blocks in the cross-correlogram (highlighted by red angle markers). Median toroidal score is indicated for each cluster with >50 putative units. Center, barcode diagram showing results of persistent cohomology analyses at P9 for the cluster highlighted in red (n = 70 units; rat #29451; longest bars in the first three dimensions: H0, H1 and H2). Grey shading indicates the longest lifetimes among 100 iterations in shuffled data (aligned to lower values of original bars). Arrows point to the four most prominent bars across all dimensions (the 2 longest bars in the H1 dimension used for toroidal decoding, neither of which is longer than shuffled data in H1 and H2. Right, toroidal rate maps for 16 representative cells. Each celĺs activity distribution is color-coded across the two H1 dimensions of the inferred torus (*ϕ*1 and *ϕ*2 H1). Brighter colors highlight the peak firing rate. Note the lack of clear tuning to a specific toroidal phase at P9. d) Same as in (c), but for a cluster from the cross-correlogram of a P10 pup (n = 121 units; rat #28801; square corners indicated in red). Arrows in the barcode plot clearly indicate the homological signature of a 2-D torus: one 0-D bar (*H*^0^), two 1-D bars (*H*^1^) and one 2-D bar (*H*^2^). The four most prominent bars across all dimensions exceed the longest lifetimes in the shuffled distribution. Right, 16 representative examples of single-cell tuning to specific positions on the toroidal manifold (*ϕ*1 and *ϕ*2 in H1). At P10, there is clear toroidal topology for the selected cluster, with individual cells each tuned to a specific toroidal phase. e) Torus-like organization at P10 uncovered by non-linear dimensionality reduction (UMAP) of the population activity in the cluster highlighted in red in panel (c). Three different view angles are shown for the same point cloud. Each dot indicates the population state at a given time point, with colors representing load on the first principal component (time bins were subsampled taking every 65th time bin, corresponding to 650–ms intervals between selected samples; see Methods). Bold black/red line represents a 5–s trajectory, showing smooth movement over the toroidal manifold. f) Toroidal ensembles appear abruptly at P10. Scatterplot shows median toroidal score as a function of age for all detected clusters with at least 50 units in MEC and PaS (P8: n=4 animals; P9: n=4 animals; P10: n=6 animals; P11: n=6 animals; P12: n=3 animals). Each dot refers to one cluster.

## Results

### Toroidal manifolds precede spatial navigation

To maximize unit yield and enable precise localization of recording sites, the majority of experiments were performed in a semi-chronic setting, in which surgery, recording, and perfusion were completed within a single day (n=15; Supplementary Table 1). In a smaller subset of animals (n=3), probes were implanted chronically, enabling repeated recordings across consecutive days. Following each session, these pups were returned to the nest to be fed by the dam. All pups, in both groups, were implanted between P8 and P12 with Neuropixels 2.0 probes targeting MEC and adjacent PaS in the right hemisphere (Fig. 1a–b). In most cases, probe trajectories traversed both structures: MEC layers 2/3 in 13 animals and PaS in 11 animals (Supplementary Table 1). Recordings were obtained on the day of implantation — and, in chronically implanted animals, on subsequent days — while the pups slept in a heated pot. Across 24 sessions (1–3 per rat), we isolated between 350 and 1,341 simultaneously recorded units (Supplementary Table 1).

To identify functional cell groups within each data set, we employed a spectral clustering approach based on intrinsic firing correlations. For each pair of neurons, we computed cross-correlations of firing rates across time lags from 0 to 300ms in 10ms bins^12,38^ (Figure 1c–d, left panels). The ratio of the minimum to maximum correlation across all lags served as a co-activity metric for each pair. Cell pairs were then clustered based on these temporal firing rate relationships, producing discrete blocks of similarly correlated neurons in the correlograms (Fig. 1c,d, left panels). The same clustering parameters were used across all sessions and animals.

To visualize the population activity of each cluster, we used a two-stage dimensionality reduction pipeline: principal component analysis (PCA) to extract the first six components, followed by Uniform Manifold Approximation and Projection (UMAP) applied to the PCA output (Fig. 1e). In parallel, we quantified the structure of the PCA-reduced point cloud using topological data analysis (Fig. 1c–d). Specifically, we used persistent cohomology to identify and measure the persistence of topological features (connected components, loops, and cavities) within the point cloud^9,11,39^ (see Methods). These features were quantified and visualized as barcode diagrams, where long-lived bars that extend well beyond values for shuffled controls indicate robust underlying structure (middle panels). A toroidal manifold has a characteristic signature consisting of one 0-dimensional feature (a single connected component), two 1-dimensional features (loops), and one 2-dimensional feature (a cavity). To ensure robustness, only clusters containing more than 50 cells were retained for further analysis.

UMAP visualizations displayed a distinct torus-like geometry emerging in one or more clusters per animal from P10 onward (Fig, 1e; Extended Data Fig. 3a–d). In persistent cohomology analyses, no evidence of toroidal topology was detected at P8–P9, whereas from P10 onward at least one cluster per animal displayed the full toroidal signature, with the lifetimes of two 1-dimensional features and one 2-dimensional feature significantly exceeding those from shuffled controls (Fig. 1c–d, middle; P < 0.01). This toroidal signature was maintained when the spiking activity of the P10 cluster was dowsampled to match firing rates at P8/P9 (two 1-dimensional features and one 2-dimensional feature consistently exceeded those from shuffled controls, P < 0.01).

To further validate the presence of toroidal structure, we projected the activity of each neuron within the identified clusters onto the angular coordinates of the two dominant 1-dimensional cohomology classes (*ϕ*1 and *ϕ*2). This projection yielded 2-dimensional firing-rate maps on the hexagonal torus, allowing us to define toroidal firing fields for individual cells. Each cell’s tuning to the toroidal manifold was quantified using a toroidal score, computed as the correlation between the cell’s firing field and a circular Gaussian on the hexagonal torus (see Methods). Consistent with the absence of toroidal signatures at the earliest ages, no reliable tuning to the inferred toroidal coordinates was observed at P8–P9 (Fig. 1c, right). Beginning at P10, however, a subset of clusters contained neurons with strong, spatially localized turning on the torus.

The highest median toroidal score per recording rose sharply — from 0.38 and 0.33 at P8–P9 to above 0.61 on all subsequent days (P10–P12; Fig. 1f). Clusters were classified as toroidal if their toroidal scores exceeded 0.60, a threshold that reliably separates toroidal from non-toroidal clusters in adult recordings (Extended Data Fig.2). Using this criterion, we found no toroidal clusters at P8 or P9 (0/4 animals each). In contrast, toroidal clusters were present in 4 of 6 animals at P10, 6/6 at P11, and 2/3 at P12 (Fig. 1f; for an example of a P11 recording, see Extended Data Fig. 3). Moreover, once toroidal tuning emerged, decoded locations produced a smooth trajectory across successive time points on both the UMAP embedding and the inferred toroidal coordinates, in contrast to the static and discontinuous dynamics observed in non-toroidal clusters (Fig. 1e; Extended Data Fig.3e, Extended Data Fig.4).

**Figure 2.**
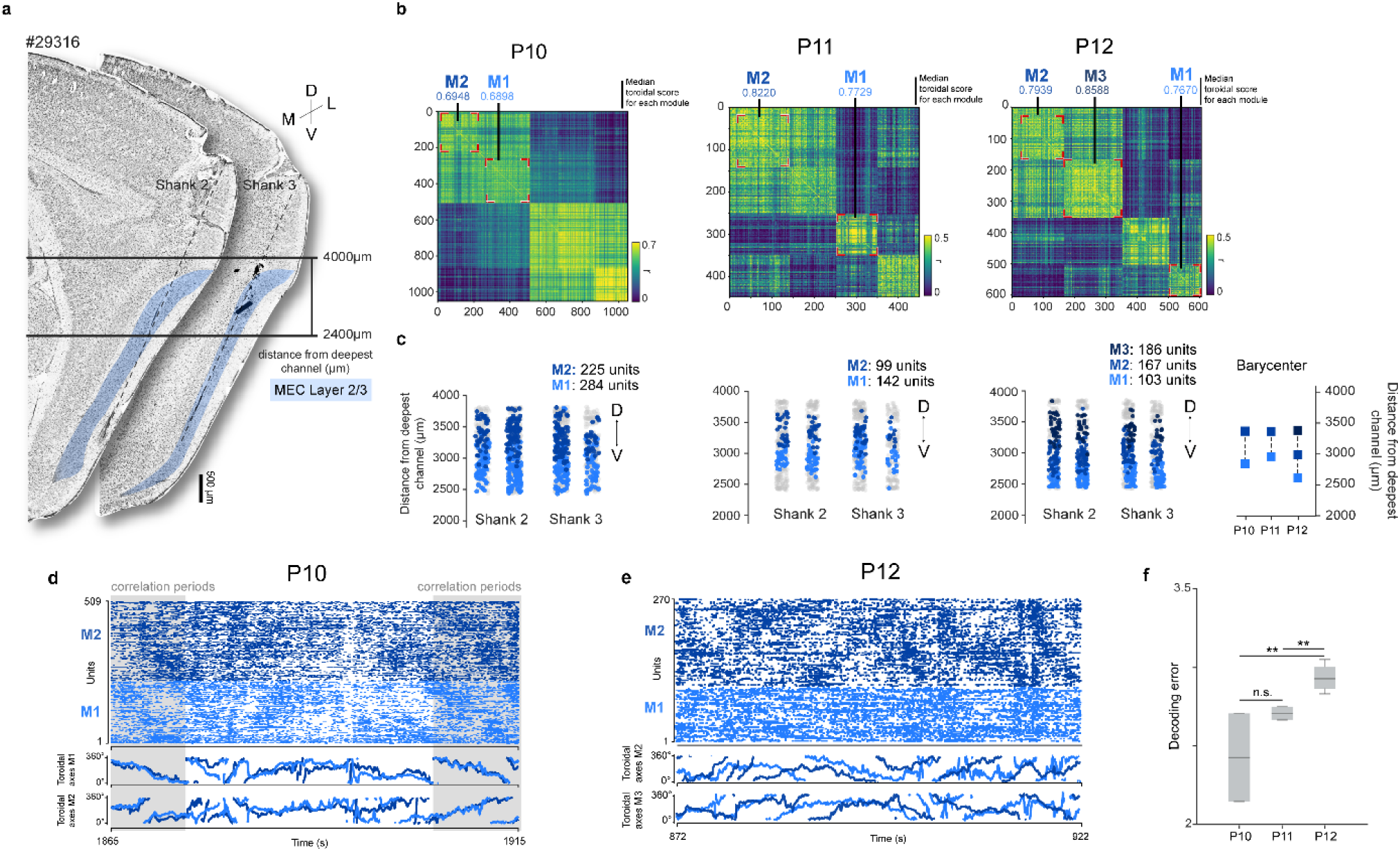
Toroidal topology has modular features from the outset. **a.** Sagittal section from a chronic rat implanted at P10 and recorded from P10 to P12, with Neuropixels 2.0 tracks highlighted by grey dotted lines. The range of recording sites across shanks 2 and 3 is marked by bold horizontal lines. Sections are organized from medial (left) to lateral (right). MEC Layer 2/3 is highlighted in light blue. **b.** Time-lagged cross-correlograms (as in Fig. 1c–d) showing a tendency for partition into toroidal modules is present already at P10 (weak distinction between M1 and M2 clusters). Modules become more distinct on P11 and P12 (Extended Data Fig. 3a), and a third grid module, M3, is additionally detected on P12. Median toroidal score for each toroidal module is reported on top. Same animal is shown across days. **c.** Recording locations of toroidal modules. Each dot represents the position of a recorded unit on the Neuropixels probe shanks. Distinct toroidal clusters are highlighted in different colors. Units that are part of non-toroidal clusters are represented by grey dots. Right panel, center of mass of each module. Note how modules are positioned successively along the dorsoventral axis. **d.** Toroidal dynamics over time at P10. Raster plot of units in two clusters classified as toroidal (modules M1 and M2, top panels) and decoded angular position (0° to 360° degrees, bottom panels) extracted via cohomological decoding over the two most prominent bars in the H1 dimension for both modules, shown as a function of time. Periods of correlation (grey shading) are interleaved by periods of decorrelation (white, non-overlapping traces). M1, light blue; M2, dark blue. **e.** Same as (d), recorded in the same animal but at P12. **f.** Mean decoding error between toroidal axes (ϕ1 and ϕ2) from one module to a second, simultaneously recorded one, at P10 (n = 4 modules), P11 (n = 4 modules), and P12 (n = 8 modules). Pairwise comparisons were performed using an exact permutation test, followed by a Benjamini-Hochberg false discovery rate (FDR) correction for multiple comparisons. P10 vs P11: effect size = 1.22, p = 0.15; P11 vs P12: effect size = 2.66, p = 0.002; P10 vs P12: effect size = 2.96, p = 0.002. Median values rose from 2.42 [2.14–2.70] on P10 to 2.70 [2.66–2.75] on P11 and 2.92 [2.86–3.00] on P12 (medians and interquartile range, IQR). Central mark indicates median; box edges represent interquartile range. The length of the whiskers indicates 1.5 times the interquartile range. **, P <= 0.01, significant after FDR correction.

These results indicate that low-dimensional toroidal manifolds arise in MEC networks by P10, well before continuous upright locomotion and prior to opening of eyes and ear canals.

### Toroidal manifolds are modular from the outset

A hallmark of the adult grid cell system is its organization into discrete modules, each defined by a characteristic grid spacing and arranged along the dorsoventral MEC axis^40,41^. We asked whether this modular organization is present at the emergence of toroidal network activity, or whether it appears only after further maturation. Comparisons across days were performed for both semi-chronic and chronic groups, with particular focus on the latter, in which recordings were obtained from the same MEC locations over multiple days (Fig. 2a).

Evidence for separation into discrete toroidal clusters was already apparent at P10 (Fig.2b, left; Extended Data Fig. 5a). At this age, 2 of the 3 chronically recorded animals had at least two simultaneously recorded toroidal clusters with toroidal scores passing the 0.60 toroidal-score threshold; Supplementary Table 1). Pairs of toroidal clusters were observed also on P11 (1/3 semichronic and 1/3 chronic animals had two toroidal cluster simultaneously recorded; Fig. 2b, middle). By P12, up to three toroidal clusters could be recorded simultaneously: 1/3 had two clusters; 1/3 had three (Fig.2b, right). As in adult MEC, whenever multiple toroidal modules were present, they were arranged topographically along the dorsoventral axis (Fig. 2c). To quantify this organization, we estimated the anatomical center of mass of each cluster based on the probe-shank positions of its constituent neurons and found an average spacing of 398 ± 26 μm (mean ± s.e.m.) between neighboring modules. Relative to non-toroidal clusters, toroidal clusters showed higher firing rates, more pronounced burst activity^11^, and tighter anatomical localization (Extended Data Fig. 5).

When examining changes in firing dynamics across all clusters (both toroidal and non-toroidal), we observed a gradual decline in overall cell-to-cell correlation (Extended Data Fig. 5a; decreasing correlation of cell pairs across postnatal days; Spearman p value = -0.357, p value = 10^−8^). Within the subset of clusters with toroidal topology, simultaneously recorded pairs of modules at P10 often displayed extensive periods of shared dynamics, during which spike traces, organized by their respective toroidal phases, followed closely aligned trajectories (Fig. 2d, top panel). By P12, such coordinated periods were markedly less evident (Fig.2e, top panel). To quantify this transition, we inferred, for each cluster, the average activity bump along the two dominant 1-dimensional cohomology classes. We then decoded the bump-activity of one module using the coordinates derived from a second, simultaneously recorded module. This provided a measure of shared dynamics: at P10, the the decoded trajectories of the two modules frequently overlapped, whereas by P12, they diverged and followed distinct paths (Fig.2d-e, bottom panels). Decoding error, defined as the mean angular difference between the decoded activity across axes *ϕ*1 and *ϕ*2, (decoding M1-from-M1 vs decoding M1-from-M2), increased significantly over the first two days after P10 (Fig. 2f; exact permutation test: P10 vs P11: effect size = 1.22, p = 0.15; P11 vs P12: effect size = 2.66, p = 0.002; P10 vs P12: effect size = 2.96, p = 0.002). Taken together, these results show that toroidal modules arise as distinct clusters from the very onset of toroidal activity. Initially, their dynamics remain partially correlated — possibly reflecting the entrainment of a common, non-spatial input — but over the following days, before the onset of exploratory behavior, these shared interactions weakened, giving rise to the fully independent modular organization characteristic of the mature grid-cell system.

### Emergence of toroidal manifolds reflects maturation of cortical network dynamics

To identify the developmental transition underlying the sudden emergence of toroidal structure at P10, we examined how population-level coordination in the MEC–PaS evolved across the surrounding postnatal days (N= 24 sessions in 18 animals). The recordings revealed a striking shift in entorhinal network dynamics around P10. At the earliest stages (P8–P9), neuronal populations displayed strongly synchronized burst patterns separated by prolonged periods of relative silence (Fig. 3a, top). At P10–P11, this pattern transitioned to a sustained, desynchronized firing regime (Fig. 3a, bottom) similar to that observed in other neocortical regions^23–25,42.^

**Figure 3.**
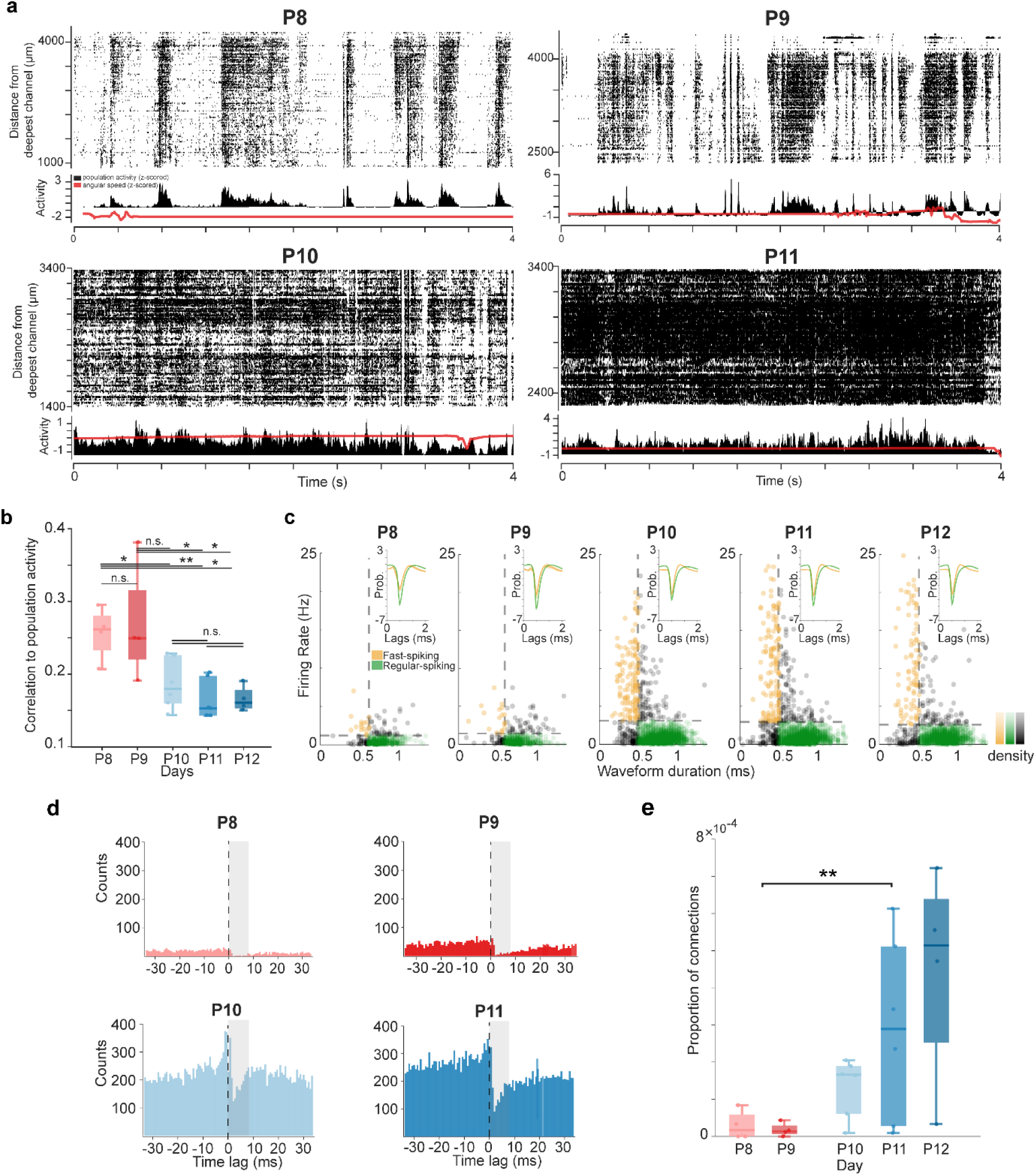
Emergence of toroidal manifolds coincides with network desynchronization and rise of peri-somatic inhibition. **a.** Raster plots of all simultaneously recorded units for representative recordings at P8–P11. Units are ordered by the recording sités distance from the deepest channel (y-axis); data are shown for a 4s time window (x-axis). Recordings are from different animals. Z-scored summed population activity (black) and angular speed (red) are plotted below. Note the shift between population activity characterized by high synchrony and abrupt shifts between periods of activity and silence on P8 and P9, and an activity regime consisting of continuous desynchronized activity on P10 and P11. **b.** Correlation of individual cell activity with population activity, from P8 to P12, at a lag of 0.1s. Data pooled across 18 animals. Lower correlation values at P10–12 indicate reduced synchrony in the neural population. Comparisons across days were performed using an exact permutation test, followed by a Benjamini-Hochberg false discovery rate (FDR) correction for multiple comparisons: p < 0.05 for all comparisons except P8 vs P9, P10 vs P11, P10 vs P12, P11 vs 12 and P9 vs P12. Additionally, over the P8–P12 period, there is a decrease in the correlation of individual cell activity with population activity (Spearman correlation, r = -0.7184, p = 0.0001). Central mark indicates median; box edges represent interquartile range. The length of the whiskers indicates 1.5 times the interquartile range. n.s., non-significant; *, P<=0.05; **, P<=0.01, significant after FDR correction. **c.** Development of fast-spinking cells in the MEC. Top right, average spike template waveforms of regular-spiking cells (RS, in orange) and fast-spiking cells (FS, in green) across developmental ages (P8 to P11). Shaded areas indicate mean ± s.e.m.. FS/RS ratio across days: P8 = 0.032; P9 = 0.031; P10 = 0.066; P11 = 0.059; P12 = 0.058. Classification of FS and RS cells was based on average firing rate (y-axis) and waveform half-width duration (x-axis) (threshold values for FS cells of 90^th^ and 10^th^ percentiles, respectively). **d.** Functional putative inhibitory connections increase sharply at P10. Panels show 4 representative cross-correlograms for spike activity of cell pairs. The decrease in firing after the activity of the pre-synaptic cells suggests inhibition of the post-synaptic cells. Grey rectangle indicates time window (0.01–8ms) used to detect putative inhibitory monosynaptic connections. **e.** Estimated probability of inhibitory connections. Probability was computed by dividing the total number of detected connections by the total number of pairs that were checked for connections. P8–9 vs P10–12: exact permutation test, effect size = 1.26, p = 0.008. Central mark indicates median; box edges represent interquartile range. The length of the whiskers indicates 1.5 times the interquartile range. **, P <= 0.01.

**Figure 4.**
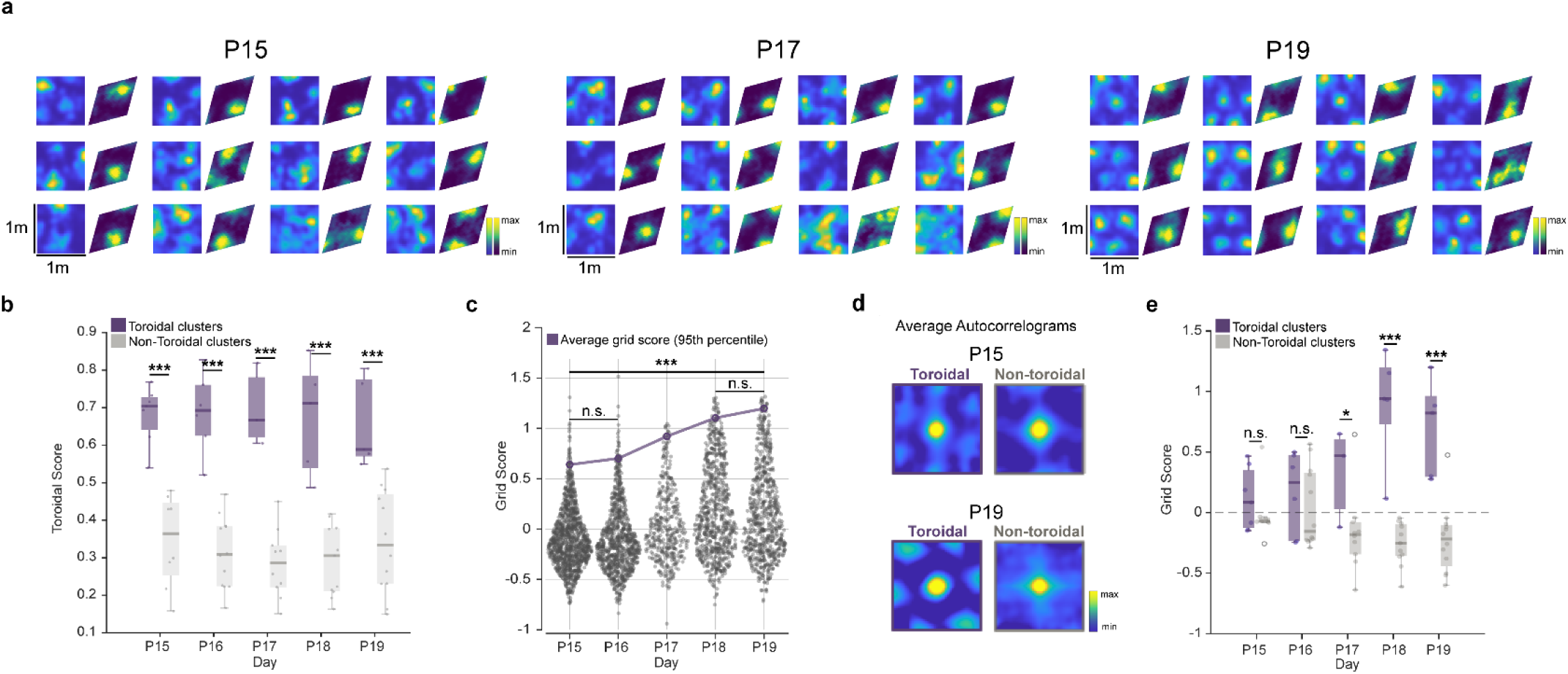
Gradual emergence of individual grid cells in toroidal manifolds. **a.** Paired spatial and toroidal rate maps from individual neurons recorded in the same rat pup on P15, P17, and P19. Different cell populations were recorded on different days. Firing rate is color-coded for each cell from minimum to maximum with respect to location in the environment (square spatial maps) or in toroidal unit space (rhomboidal toroidal maps). For each day, the 12 cells with the highest grid scores are reported, sorted from left to right according to their grid scores. Note progressive emergence of spatially periodic firing fields across days, with canonical adult-like grid patterns present by P19. **b.** Box plot showing the median toroidal score for all detected toroidal and non-toroidal clusters (purple and grey, respectively) from the onset of exploration on P15 until recordings ended on P19. Toroidal clusters have consistently higher toroidal scores than non-toroidal clusters. Exact permutation test, effect sizes in order: 3.25, 2.79, 4.3, 3.3, 2.3. All comparisons p < 10^−3^. Central mark indicates median; box edges represent interquartile range. The length of the whiskers indicates 1.5 times the interquartile range. Note stability of toroidal scores across days of recording. ***, P <= 0.001. **c.** Spatial regularity increases over days. Plot shows distribution of grid scores for all cells of all toroidal clusters in all animals (P15, n=3; P16 and P17, n=4; P18 and P19, n = 5). Each dot represents one unit (from 701 to 1,289 per day). Grid scores increase from P15 to P19 (Kruskal-Wallis test, χ^2^= 349.60, p = 10^−74^; Dunn’s Post-Hoc tests: p > 0.05 for P15 vs P16 and P18 vs P19 comparisons; all other comparisons: p < 10^−3^). Purple circles and line show 95^th^ percentile values for all cells across days. **d.** Representative examples of an average spatial autocorrelogram of all neurons in toroidal versus non-toroidal clusters on P15 (top) and P19 (bottom). Correlation values are color-coded from minimum to maximum. A clear hexagonal pattern, a hallmark of mature grid cells, is evident in the toroidal cluster at P19. **e.** Box plot showing distribution of grid scores for average autocorrelograms (as in d) in toroidal vs non-toroidal clusters (purple and grey). Data are from all animals (same as in b–c). Central mark indicates median; box edges represent interquartile range. The length of the whiskers indicates 1.5 times the interquartile range. Outliers are indicated by empty circles. n.s., non-significant; *, P <= 0.05; ***, P <= 0.001.

To quantify this transition in MEC–PaS dynamics, we computed, for each age group, the correlation between individual unit activity and the global population activity, estimated by summing the spike counts of all simultaneously recorded neurons within discrete time bins. Using 100ms time bins, we observed that median correlation values declined sharply from 0.26 (interquartile range, 0.23–0.28) at P8 and 0.24 (0.22–0.31) at P9 to 0.17 (0.15–0.22) at P10, 0.15 (0.14–0.19) at P11, and 0.16 (0.15–0.17) at P12 (decreasing correlation of individual cells to population activity across postnatal days; Spearman correlation, r = -0.7184, p = 0.0001), indicating a robust decrease in network synchrony starting on P10 (Fig. 3b). The reduction persisted when the number of units was matched across days (50 randomly sampled units for 200 iterations; Spearman correlation, r = -0.7609, p = 0.00003). The effect was preserved across bin sizes from 50ms to 500ms (Extended Data Fig 6a) and in pairwise correlations (Extended Data Fig. 6b). These observations point to a coordinated shift in population dynamics coincident with the emergence of toroidal topology.

To identify physiological mechanisms accompanying this desynchronization, we examined developmental changes in local excitatory-inhibitory balance, focusing on the emergence of perisomatic inhibition, a key regulator of timing and synchrony in cortico-hippocampal networks^42–44^. Among well-isolated putative units (see Methods), the proportion of fast-spiking units increased from 3% at P8–P9 to 6% at P10–P12 (Fig. 3c). This increase was expressed by a higher proportion of neurons with narrow spike waveforms (10^th^ percentile waveform widths of all cells decreased from 0.63ms at P8 and 0.60ms at P9 to 0.50 ms on each day between P10–P12) and by an increase in average firing rates across the whole recorded population (90^th^ percentile values of 1.2 Hz at P8, 1.4 Hz at P9 and from 2.5 Hz to 3.10 Hz between P10 and P12) (Fig. 3c; Extended Data Fig. 7a). Together, these observations suggest a rise in interneurons recruitment at P10.

We next asked whether the growth in fast-spiking activity was matched by a corresponding rise in inhibitory synaptic connectivity. To test this, we analyzed spike train cross-correlograms (CCGs)^29,45,46^ for all recorded cell pairs across 18 animals to identify putative monosynaptic inhibitory connections. Cell pairs were classified as putatively inhibitory when the firing probability of the postsynaptic neuron showed a reliable suppression for at least 2 ms within the initial 0.01-8 ms after the presynaptic spike (see Methods, Fig. 3d; Extended Data Fig. 7c). The fraction of such pairs increased severalfold at P10–P12, from between 0.0038 to 0.0084 permille at P8–P9 to 0.0471 to 0.0940 permille at P10–P12 (Fig. 3d and Fig. 3e; exact permutation test, effect size = -1.26, p = 0.008). At P10–P12, most pre-synaptic cells in these connections exhibited short-latency waveforms and high firing rates (Extended Data Fig. 7), consistent with an interneuron identity^47^ and indicating that the rise in interneuron activity is accompanied by increased functional inhibitory connectivity.

Collectively, these findings suggest that the emergence of toroidal manifolds in the MEC coincides with a sharp desynchronization of cortical network activity and the establishment of functional inhibitory circuitry. The sharp transition in network dynamics at P10 mirrors the well-documented decorrelation observed across multiple cortical regions between P8 and P12 ^23,24,42^ — a shift widely interpreted as a hallmark of emerging functional inhibition and recurrent excitatory-inhibitory connectivity^43^. The maturation of these circuit motifs in MEC and other cortices may be essential for the formation of continuous attractor dynamics thought to underlie the toroidal topology of grid cells^4,17,48,49^.

### Alignment between toroidal and physical space

Having established that toroidal population dynamics in MEC emerge before the onset of navigational experience, we next asked when these preconfigured manifolds become aligned with external spatial coordinates. To address this, we performed chronic recordings of MEC activity during foraging in an open field, starting on P15 when the animals begin to leave the nest^50^, and continuing until P19, yielding between 701 and 1,289 simultaneously recorded units in 6 animals. Time-lagged crosscorrelation and persistent-cohomology analyses were applied using the same parameters as at younger ages (Fig. 1c,d; see Methods).

Clusters identified as toroidal through topological data analysis maintained high toroidal scores throughout the P15–P19 recording window (Fig.4a–b). In contrast, the spatial tuning of cells in these clusters was initially weak, and clear grid structure emerged only gradually over subsequent days (Fig. 4a). To quantify this developmental refinement, we computed grid scores (in physical space) for all units within toroidal clusters. As expected, given that these clusters likely include non-grid cells whose activity is correlated with grid cells, the distribution of grid scores remained broad (Fig, 4c). Nevertheless, the upper tail of the grid-score distribution (95^th^ percentile) rose sharply from P15 onwards (Fig. 4c, 95^th^ percentile values increased from 0.682 ± 0.024 on P15 to 1.188 ± 0.012 on P19, means of percentile values ± s.e.m. across animals; Kruskal-Wallis test: χ^2^ = 129.4, p = 10^−74^), a value comparable to grid scores of grid cells in adult animals^6,40^. No comparable increase in grid scores was observed in non-toroidal clusters (Extended Data Fig. 8a).

To further characterize the emergence of spatial periodicity, we examined average autocorrelograms of firing fields across the P15–P19 recording window. Consistent with the development in grid scores, toroidal clusters showed progressively clearer hexagonal patterns across days, whereas non-toroidal clusters failed to develop hexagonal structure (Fig. 4d–e, Extended Data Fig. 8b). From P17 onwards, grid scores of toroidal clusters were significantly higher than those of non-toroidal clusters (exact permutation test at P17: effect size = 1.42, p = 0.04; P18: effect size = 3.94, p = 0.0002; P19: effect size = 2.87, p = 0.0008). Grid spacing in toroidal clusters was centered around peaks at 43.1 cm and 74.1cm (Extended Data Fig. 8c), comparable to adult values for grid modules in dorsal MEC^40^.

These results demonstrate that while toroidal manifolds in MEC are internally generated, their alignment with external spatial coordinates emerges gradually along with active navigational experience.

### Ring manifolds precede toroidal topology

We finally asked whether the emergence of toroidal manifolds is preceded, or accompanied, by simpler topological structures such as rings. Ring-like topology is thought to be fundamental to the operation of head direction^13,14,16,51^ and other direction-tuned neurons^29^, and in mammals, PaS cell populations with ring topology may provide directional input needed to drive the activity bump on the torus in MEC grid cells^4,8^. To test whether ring-like manifolds appear as early as, or earlier, than toroidal structures, we screened time-lagged crosscorrelograms of P8–P12 MEC-PaS recordings (Fig. 5a–b; Extended Data Fig. 1), using the same criteria for separation of clusters as in the toroidal analyses (See Methods).

**Figure 5.**
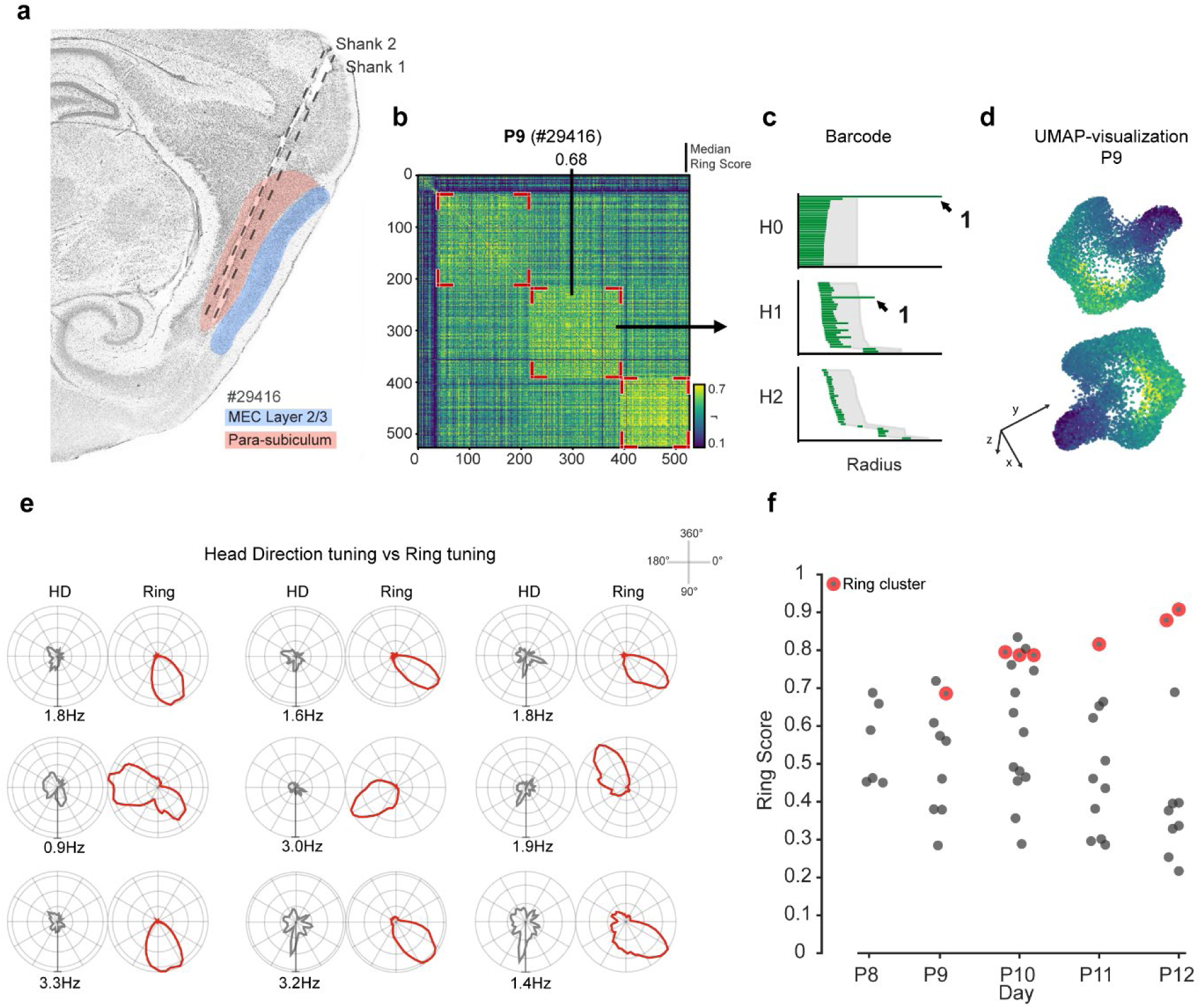
Ring manifolds are present by P9. **a)** Sagittal section from a rat implanted with a Neuropixels probe at P9. Estimated probe tracks are highlighted for two shanks by dotted lines. The shanks are clearly positioned in parasubiculum (PaS). MEC Layer 2/3 is highlighted in blue, PaS in red. **b)** Time-lagged crosscorrelogram showing correlations between all pairs of PaS neurons recorded simultaneously in one session at P9 in the rat shown in **a**. Symbols as in Fig. 1c–d, with identified clusters highlighted by red angle markers. The cluster indicated by black arrows has 173 cells and a median ring score of 0.68 (for reference, see Extended Data Fig. 9 and 10). c) Barcode diagram showing results of persistent cohomology analyses at P9 for the cluster highlighted by black arrows in **b**. Grey shading indicates the longest lifetimes among 100 iterations in shuffled data (aligned to lower values of original bars). Black arrows point to the two most prominent bars across all dimensions. Note that only one bar in H1 is longer than shuffled data, which is indicative of ring topology. d) Ring-like organization uncovered by non-linear dimensionality reduction (UMAP) of the population activity in the cluster highlighted in panel (b). Two different view angles are shown for same point cloud. Each dot indicates the population state at a given time point, with colors representing the first principal component. Time bins were subsampled taking every 15th time bin, corresponding to 150–ms intervals between selected samples (see Methods). e) Polar tuning profiles of 9 representative ring-cluster units from the ring-topology cluster recorded at P9 (same cluster as in panels b–d). Each pair compares the unit’s tuning to the external head direction (left, grey) versus the inferred position on the internally organized ring (right, red). The radial axis represents firing rate (Hz). Plots are peak-scaled for each unit; the value below each pair indicates the maximum firing rate corresponding to the outermost radial ring. The angular axis represents direction (0°–360°) plotted clockwise, with 0° aligned to the vertical top axis. f) Scatterplot showing median ring score as a function of age for clusters with at least 50 units in MEC–PaS (P8: n=4 animals; P9: n=4 animals; P10: n=6 animals; P11: n=6 animals; P12: n=3 animals). Each dot refers to one cluster. Clusters previously classified as toroidal have been removed. Note that multiple clusters from P8 exhibit high median ring scores, although clusters meeting all criteria for classification as ring-tuned are observed only from P9 onward.

Evidence of ring topology was present by P9. In one P9 cluster (1 animal out of 4), we identified a single, persistent first homology class (H1) with a lifetime significantly longer than that of shuffled data, confirming the presence of a 1D ring structure (Fig. 5c). Dimensionality reduction using PCA followed by UMAP likewise revealed a distinct ring-like manifold (Fig. 5d) and individual neurons displayed sharp tuning to the ring manifold (Fig. 5e). Similar tuning was also present in some neurons at P8 (Extended Data Fig. 9). As expected at ages when eyes and ear canals remain closed, neurons tuned to the ring manifold at P8 and P9 did not yet show tuning to the animal’s actual head direction (Fig. 5e and Extended Data Fig. 9, grey plots), consistent with prior findings that head direction signals align with environmental cues only from P11–P12 onward^20,33,34^.

To quantify tuning to the ring, we applied two complementary criteria across all clusters, in addition to the presence of a single persistent first homology class (H1). First, we computed for each neuron a ring score, defined as the correlation between an idealized 1D Gaussian centered at the neuron’s preferred direction and its firing rate distribution on the ring (mirroring the approach to computing toroidal scores in 2D). Clusters with ring scores below 0.63 (corresponding to the 25th percentile of the ring score distribution during similar behavior in adult head direction cells; Extended Data Fig. 2c), were filtered out. Second, we used a “largest gap” heuristic^52^ which selects clusters with the largest separation between the first and second circular coordinates in H1 (excluding toroidal clusters, which also exhibit high ring scores). These three criteria reliably identified ring-like topological structures in clusters comprised of head direction-tuned cells in adult MEC–PaS recordings obtained during comparable behavioral conditions (in sleep) (Extended Data Fig. 2c).

Applying these criteria to the developmental dataset, we recovered clusters meeting all three requirements as early as P9 (P8: 0 clusters; P9: 1 cluster; P10: 3 clusters; P11: 1 cluster; P12: 2 clusters). High median ring scores were already observed at P8 in 3 of 12 clusters (Fig. 5f, Extended Data Fig. 9), but these clusters satisfied only one of the classification criteria. Most ring clusters were in PaS (5 out of 7 clusters) from P9 to P12; the two remaining were in MEC L2/3 at P10 (Extended Data Fig. 10). The two ring clusters in MEC appeared before detectable toroidal topology in these animals, raising the possibility that ring manifolds in MEC serve as precursors to toroidal manifolds.

## Discussion

This study demonstrates that toroidal manifolds in MEC population activity emerge well before the onset of spatial navigation. By referencing neural dynamics to the population itself rather than to external spatial frames^35^, we were able to identify toroidal manifolds at P10, prior to eye and ear opening and the onset of upright navigation, at a time when movement is largely restricted to pivoting on the abdomen within the nest^21^. Even at this early stage, the toroidal manifolds exhibit signs of modular organization^53^. Their emergence coincides with a major shift in cortical dynamics: a transition from synchronized, burst-like activity to desynchronized and decorrelated patterns, accompanied by a notable increase in inhibitory connectivity. A second developmental phase begins around P15–P16, when animals first explore beyond the nest. From this point onward, the toroidal manifold activity progressively aligns with the external environment, culminating in periodic, grid-like firing patterns anchored to physical space by P19. Ring-shaped manifolds, associated with directionally tuned neurons, emerge even earlier, (at P9 or before), although their regularity continues to improve with age. Together, these observations suggest that low-dimensional manifolds with torus and ring topology arise through intrinsic self-organizing mechanisms that are only subsequently mapped onto the external world through experience.

These findings provide mechanistic insight into the developmental emergence of continuous attractor networks, long proposed to underlie grid cell dynamics^4,15,48,49^. In such models, grid cells are arranged on a toroidal manifold, where each cell’s position on the torus reflects its firing location in physical space, and population activity moves smoothly in correspondence with the animal’s movement. The present findings illuminate the maturational origins of this attractor-network structure, pointing to a process driven primarily by intrinsic network architecture, including recurrent connectivity. At the time when the first toroidal manifolds appear, the MEC undergoes a transition from globally synchronized activity to a more desynchronized and decorrelated state — a shift observed widely across cortical regions during the second postnatal week^23,54^. The maturation of local inhibitory circuits^44^ likely contributes to this transition by disrupting global synchrony and reducing reliance on bottom-up sensory input, thereby enabling the self-sustained recurrent dynamics required for continuous attractor states.

While toroidal manifolds emerge alongside with intrinsic network maturation, ring-shaped manifolds appear to follow a distinct developmental trajectory. Signatures of ring manifolds are evident by P9, with some individual cells displaying directional tuning as early as P8, preceding the appearance of toroidal structure at P10. This early onset raises the possiblity that ring-like architectures are shaped by subcortical inputs, such as projections from the anterior thalamus and the mammillary bodies, whose internal dynamics mature earlier than those of the cortex^55,56^. The early ring attractors in MEC may act as teaching signals that help establish phase relationships and periodicity in nascent toroidal manifolds around P10.

Finally, we showed that toroidal manifolds align with external spatial coordinates only after several days of exploratory experience. This delayed anchoring is consistent with earlier single-cell studies in juvenile rats, which observed only weakly periodic grid cell activity during initial exploration (P15–P16) and full spatial periodicity only after several additional days of experience^18,19^. The idea that experience is required to stabilize internal representations is further supported by work showing that depriving animals of geometric cues, such as rearing them without sharp boundaries, delays the maturation of stable spatial single-cell tuning^57,58^. Our findings leave open the possibility that non-spatial sensory experience contributes to the formation of a toroidal map before or around P10. Self-generated motion signals from spontaneous twitches and undirected head movements may provide activity-dependent teaching signals^59–61^ that help shape the emerging toroidal structure. Together with inputs from earlier-maturing networks with ring topology, these intrinsic signals may support the initial organization of the manifold before it becomes anchored to structured sensory and spatial inputs during active exploration.

These results motivate a two-stage model for the development of spatial mapping in the MEC. The first stage, occurring before locomotion and before the eyes and ear canals open, is intrinsically driven: low-dimensional manifolds arise from synaptic connectivity and emerging network topology^62^. During this period, the MEC–PaS circuit develops the recurrent architecture and excitation-inhibition balance required for continuous attractor dynamics. This maturation follows a structured progression — beginning with the emergence of ring-like manifolds in PaS. This is followed by the appearance of toroidal manifolds and early modular organization in MEC subpopulations, and culminating in the refinement of these modules as they disentangle from shared upstream inputs and likely gain the capacity to incorporate velocity signals. The second stage covers the subsequent postnatal week, beginning with the onset of active exploration. Experience-dependent calibration and richer sensory inputs now allow the preconfigured networks, grid cells and head direction cells alike, to anchor their activity to external spatial coordinates. This anchoring unfolds slowly, likely reflecting the time required to coordinate immature motor systems with evolving sensory feedback^4^. Through repeated action-based experience, the intrinsic network gradually learns to integrate multimodal inputs into coherent and flexible spatial representations, ultimately enabling the toroidal activity bump to track the animal’s physical location in real time.

All together, these findings suggest that low-dimensional manifolds, such as rings and tori, are fundamental organizational structures used by the MEC–PaS circuit as soon as intrinsically-driven dynamics are present. The results support the general idea that cognitive operations may rely on evolutionary predetermined neural architectures that are later calibrated and anchored to the external world through experience. Such architectural priors may likewise be leveraged in artificial systems to efficiently parse and navigate the world in ways analogous to biological systems^63^.

## Methods

### Subjects

The data were obtained from 23 Long Evans rat pups between postnatal days P8 and P19 (20 males and 3 female; between 20g and 51g at the time of implantation). Sex, age, and weight for each individual rat is provided in Supplementary Table 1.

All animals were bred in-house. P0 was defined as the first day when a litter was observed; pregnant dams were checked multiple times per day between 7 am and 9 pm. All animals were on a reversed day/night cycle, with room lights turned on at 7 am and off at 7 pm and no lights turned on between 7 pm and 7 am. A maximum of 2 rats were implanted in each litter. If pups were reintroduced to the litter for chronic experiments, only one animal per litter was implanted. Litter sizes were reduced to a maximum of 6 after surgery; surplus pups were either euthanized or passed on to adoptive dams. Experiments were approved by the Norwegian Food Safety Authority (FOTS ID 18011 and 29893) and conducted in accordance with the Norwegian Animal Welfare Act and the European Convention for the Protection of Vertebrate Animals used for Experimental and Other Scientific Purposes.

### Surgery and electrode implantation

All animals were implanted with Neuropixels silicon probes (prototype 2.0 multi-shank probes) targeting MEC layers 2 and 3 (L2/3) and parasubiculum (PaS)^11,36^. Probes were always implanted in the right hemisphere at different ML coordinates according to ages: 4.0–4.5 mm lateral to the midline for implants between P15 and P19, and 3.80–4.00 lateral to the midline for implants between P8 and P12. All implants were positioned 0.0–0.2 mm anterior to the transverse sinus, at an angle of 12-22° in the sagittal plane, with the tip of the probe pointing in the anterior direction. Probes were lowered to a depth of 4,000–5,000 μm, or until one of the shanks hit the cranial bone. Final implantation depth was noted down. The implants were secured with dental cement and with a custom made 3D printed head cap (in black resin), to protect the implant once the pup was reintroduced to the dam and litter. A jeweller’s screw was placed in the skull above the cerebellum and was connected to the probe ground and external reference pads with an insulated silver wire. The surgical procedure always lasted less than 3h, to facilitate reintroduction of the pup in the litter (in case of chronic experiments).To prevent rejection, dams were habituated in advance to the 3D-printed caps used to protect the implants. Analgesia (meloxicam) was administered during the recovery period according to guidelines. Recordings began when rats had recovered from anesthesia at least 1 hour after surgery.

### Electrophysiological recordings

The instrumentation and protocols employed here were similar to those described in previous work from the lab^11^. Briefly, the probe’s integrated circuitry was responsible for amplifying, filtering, and digitizing neural signals at 30 kHz. The probes used a gain of 80 and a 0.005–10 kHz filter. Signals were multiplexed and transmitted to the recording system through a tether cable. We utilized SpikeGLX (https://billkarsh.github.io/SpikeGLX/) for both probe configuration and acquisition.

To monitor 3D head orientation and position, we implemented a motion-capture setup comprising OptiTrack Flex 13 cameras, Motive software, and retroreflective markers attached to the implant. The 3D coordinates were projected onto the horizontal plane to calculate head-direction azimuth and 2D position.

Data stream timestamps were synchronized using previously described methods^11^. An Arduino microcontroller generated randomized digital pulse sequences that were were routed as direct TTL inputs to the Neuropixels acquisition system, while simultaneously being signaled to the video camera and OptiTrack system via infrared LEDs positioned within the recording perimeter of the test environment.

During experiments the layout of recording sites on the probe shanks was chosen based on spike activity and implantation depth at surgery (between 4 and 5 mm from the brain surface). The final recording location of individual electrode contacts was confirmed ex-vivo by aligning a map of the probe shanks to histological sections, using the probe tip as a reference point.

### Experimental procedures and behavior

Following the surgical procedure, the pups were allowed to recover in a heated chamber for at least 1h. They were then transferred to the experimental room and placed on a towel in a large flower pot (33cm diameter) on a pedestal to start experiments. The flowerpot was constantly heated with hand warmer pockets. For P8–P12 animals, the experimental session consisted of a 2h sleep session in the heated flowerpot. For animals implanted at P14–P18, the experiment consisted of a foraging session in a 1 × 1 m open field arena, with dark green flooring and enclosed by black walls of 50 cm height. Multiple cue card were affixed to the walls of the arena. A dim light in the left corner of the room was left on during the experiment. All experiments were performed in the same recording room, utilizing the same distal and proximal cues. The open field session typically lasted for 60 min. Food crumbs (small cookie crumbs) were randomly scattered in the arena. In semi-chronic animals, the experimental session was followed by perfusion to extract and precisely localize the probe implant position. In chronic animals, pups were returned to the dam and littermates. To facilitate the reintroduction of the pups we always removed the mother from the cage and reintroduced the siblings. To facilitate the reintroduction of the pups we always removed the dam from the cage and reintroduced the siblings. None of the animals were ever food or water deprived.

### Spike sorting and single-unit selection

Spike sorting was performed using KiloSort 2.5^36^, using custom modifications for high cell-density recordings in MEC–PaS as previously described in^11^. Units were first classified as multi-unit activity (‘MUA’) based on a >10% estimated contamination rate from spikes from other units, computed from actual vs expected refractory period violations from the autocorrelogram. Units with large waveform spatial footprints (> 100 μm) were also removed, as described previously^29^. Units classified as ‘MUA’, were removed from monosynaptic connectivity analyses. Burst events were defined as consecutive spikes separated by an inter-spike interval (ISI) of less than 7 ms. The burst proportion for each neuron was calculated as the ratio of burst events to the total number of firing events, which included both burst events and single spikes (ISI >7ms).

### Clustering

During postnatal days P8–P12, pups exhibit a limited repertoire of behaviors and spend most of their time asleep^22,64^, precluding the use of behavioral variables variables to identify functionally defined cell types such as grid cells and head direction cells. To identify functional cell groups within each data set of the present work, we employed a spectral clustering approach solely based on cell-to-cell firing correlations, using the raw spike-time data obtained from automated spike sorting. To ensure robust clustering, we excluded units which did not meet the following three criteria:

1. Firing rate: Units with a mean firing rate < 0.2Hz or > 4Hz were excluded.
2. Population coupling: To exclude units driven primarily by global population fluctuations, we calculated the Pearson correlation (*r*) between each unit’s smoothed firing rate (100ms Gaussian kernel) and the population mean activity. We fitted a bimodal distribution to the resulting *r*-values and defined an upper exclusion threshold at the peak of the lower mode plus two standard deviations.
3. Sparseness: we calculated the standard deviation (σi) of the binned spike counts for each unit. unit. Units were excluded if more than 50% time bins had a spike count *c* < σi.

Following exclusion (on average 40% of simultaneously recorded units in a session), clusters were identified based on the temporal co-activity relationships within the remaining population, as described in a previous study^12^. We first computed the cross-correlation function, *c*ij(*ττ*), between the smoothed firing rates (sampled at 100ms intervals and using a gaussian smoothing window with σ = 100ms) for all pairs of neurons over lags *τ* in [0, 3*s*].

To capture cluster membership, we defined, for each cell pair, a scalar similarity metric, *Mij*, as the ratio between the minimum and maximum cross-correlation values over the lag window:

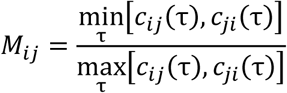

We then constructed a similarity matrix, *S*, where each entry, *Sij*, is the Pearson correlation coefficient between the *i*-th and *j*-th rows of *M*. This measure quantifies how similarly two neurons relate to the rest of the population. *S* was row-normalized to produce a stochastic matrix, *X*:

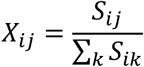

We applied the spectral clustering algorithm described in^65^, using code from https://github.com/smpcole/clustered-markov, to the Laplacian (*L = I – X*) of this matrix. The optimal threshold determining cluster count was defined by the midpoint between the “elbow” of the 40 first singular values (found using the *KneeLocator* python-package) and the subsequent singular value. Clusters containing fewer than 50 units were discarded.

### Preprocessing of neural data

Following clustering, we applied preprocessing steps adapted from previous work^11,12^. Spike counts were binned into 10 ms intervals and smoothed using single exponential smoothing (via the SimpleExpSmoothing class in *statsmodels*), followed by a square-root transformation. The smoothing parameter *α* was set to 0.1 for toroidal analyses and 0.05 for ring analyses, reflecting the slower dynamics of head-direction cells compared to grid cells. To mitigate temporal autocorrelation bias between consecutive time steps and to standardize dataset size, we uniformly subsampled the data to 120,000 time points per session. The data were then speed-filtered based on state and age: for awake sessions (>= P15), periods of immobility (< 2.5 cm/s) were excluded; conversely, for sleep sessions (< P15), periods of movement (> 0.1 cm/s) were removed. We further filtered for high-activity states by retaining only the top 50% of time points with the highest total population activity. To remove noise and isolate the underlying manifold of the population activity, we performed dimensionality reduction on the resulting *N*-dimensional point cloud (where *N* is the number of neurons). The activity of each neuron was first *Z*-scored to normalize contributions, and the first 6 Principal Components (PCA) of the population activity was retained. These components were whitened by their corresponding eigenvalue and normalized.

Finally, the resulting 60,000 dimension-reduced data points were further reduced to 1,600 points to allow for computing persistent cohomology. This was achieved using the two-step downsampling technique described previously^12^. (1) “Radial” downsampling was first applied: iteratively selecting the closest point with a minimum Euclidean distance threshold of 0.15 from the current set of chosen points. (2) “Fuzzy” downsampling, using a Gaussian kernel density (with a neighborhood of 5% of the points to define kernel width) to iteratively select the point of highest density, subtracting the chosen point’s contribution from the density function in the next iteration.

### Persistent cohomology

We characterized the topological structure of neural activity using persistent (co-)homology framework, following the workflow of previous papers describing the toroidal topology of grid cells activity^11,12^, and the ring topology of head direction cell activity^10,66^. Briefly, the algorithm approximates the underlying topology by expanding balls centered on each data point. As the radius increases, the union of these balls forms a space containing holes of various dimensions. The interval of radii for which a hole exists is recorded as its ‘lifetime’ and the union of these intervals constitutes the barcode.

To validate topological features, we generated shuffled distributions by independently rolling the firing rate arrays a random length between 0 and the length of the session. The same preprocessing and persistence analysis were performed on the shifted spike trains as for the unshuffled data. We repeated the analysis on 100 such shuffled datasets to determine the maximum feature lifetime likely to occur by chance. This maximum value defined the significance threshold for each dimension.

The software package Ripser^67,68^ was used for all computations of persistent cohomology.

### Cohomological decoding and toroidal rate maps

We obtained toroidal coordinates using the same cohomological decoding framework described by^69^ and previously applied in^11,12^. Briefly, this method uses the harmonic cocycle representative of the 1-dimensional holes found using persistent cohomology to obtain a function mapping each point in the point cloud to a representative angle on the detected circle. For toroidal analyses, we decorrelated each pair of circular coordinates using the algorithm introduced by^70^. This gave toroidal coordinates for the 1600 points in the downsampled point cloud. To obtain coordinates for the population vectors excluded in the downsampling process, we extrapolated the coordinates. We first computed probability distributions for each neuron’s activity on each detected circle and for each remaining vector, found the weighted mass cented on the torus, weighing each distribution by the respective neuron’s activity in the given population vector and summing the distributions across all units in the cluster. Toroidal ratemaps were constructed by binning the toroidal surface into a square grid of 10° × 10° bins and computing the average activity in each position bin. To spatially smooth the toroidal rate map we convolved with a 2-D gaussian filter of 1-bin kernel width.

### Torus and ring scores

In adult rats, grid cells have single confined tuning fields on the underlying torus. To quantify how well neurons were tuned to the decoded torus, we computed a toroidal score by correlating each toroidal rate map (computed using 10 × 10 bins) with that of an idealized Gaussian bump (σ = 0.1). The Gaussian profile was defined over 1600 points randomly sampled from a hexagonal torus and was centered at the same mass center as the neuron under analysis. A hexagonal torus is characterized by three axes oriented 60 degrees apart. Because only two of these are linearly independent and we do not know a priori which pair of axes we find using persistent cohomology, we computed the toroidal score assuming both possible orientations (60 and 120 degrees) and choosing the orientation based on the highest median score across the population. We additionally optimized the Gaussian width σ to the hexagonal torus by sampling 50 values between 0.01 and 0.5 radians.To calculate 1-D ring scores, we correlated each ring tuning map (computed using 10 bins) with an idealized 1-D Gaussian bump (σ = 0.1) on a circle. The 1-D Gaussian profile was defined over points randomly sampled across the circular manifold and centered at the same mass center as the selected neuron.

The classification of toroidal and ring clusters in pups was compared to values obtained from data in adult animals, using a published dataset^29^.The following animals from this dataset, all with a clear grid cell population, were included: 28229_2, 25691_1, 25843_2, 26034_3, 27765_3, 28304_2. The recordings were analyzed using the same pipeline as described above, separating MEC and hippocampal recordings. We used the study’s MUA classification to exclude noisy cells. For grid cell analyses, we used awake open field sessions, and the clustering threshold was manually set to: 0.98, 0.96, 0.97, 0.93, 0.75, 0.98 for each animal (in the same ordering as listed above). Head direction cells were identified using the mutual information score between the firing rate activity of each cell and the simultaneously recorded head direction, requiring scores above 1 bit and excluding cells classified as a grid cell in the study. Ring scores were computed during sleep sessions, to match experimental conditions in pups (P8–12).

### Visualization of toroidal and ring manifolds

For each cluster of neurons forming a toroidal module, spike times of co-recorded cells were binned at 10 ms resolution and convolved with a Gaussian kernel (σ = 50 ms). In pups between P8 and P12, behavior is dominated by rest states, and locomotor speed does not reliably separate periods in which toroidal dynamics are expressed. Instead, brief movements or twitches can introduce transient motion-related artifacts, therefore time bins with speed above a permissive threshold were excluded. To further identify moments in which toroidal dynamics are likely to be expressed, we retained only time bins with sufficient overall neural activity, defined as those exceeding the mean population firing rate (Z-scored population rate > 0). Low-activity bins were removed because they are unlikely to maintain enough momentum to drive coherent toroidal dynamics. Smoothed firing rates were z-scored per neuron to normalize across firing-rate differences. For computational tractability, time bins were downsampled by selecting every 65th bin (equivalent to 650 ms between samples). The resulting population activity matrix was treated with time bins as observations and neurons as variables, and principal component analysis was applied, retaining the first six components. Uniform Manifold Approximation and Projection (UMAP) was then run on these components using the following hyperparameters: ‘n_dims’=3, ‘metric’=‘cosine’, ‘n_neighbours’=5000, ‘min_dist’=0.8 and ‘init’=‘spectral’. The continuous trajectories were reconstructed from discrete timepoints in the UMAP space using spline interpolation. For visualization of the ring manifold at P9, we applied an activity filter to reconstruct the topology from discontinuous data: analyses were restricted to active epochs by excluding time bins where the population firing rate dropped below 5% of the maximum. Time bins were downsampled by selecting every 15th bin (equivalent to 150 ms between samples). The remaining valid time points were then concatenated for dimensionality reduction.

### Waveform information and classification

During spike sorting with Kilosort, each unit is assigned with a 2 ms spike waveform template on each recording channel. For each unit, the representative waveform was defined as the temporal profile recorded on the channel yielding the maximum absolute amplitude. Only units manually curated and labeled as ‘good’ were included in the analysis. To ensure high temporal precision for metric calculation, the original waveforms (sampling rate: 30 kHz; 61 samples) were upsampled by a factor of 10 using linear interpolation. Peak-to-trough duration was defined as the time interval between the global minimum (trough) and the subsequent local maximum (peak) of the waveform. Rather than applying fixed cutoffs, we established dynamic thresholds based on the population statistics of each specific day. Fast-spiking units were defined as units exhibiting a narrow waveform (duration < the daily 10th percentile) and a high firing rate (rate > the daily 90th percentile). . Regular-spiking cells were defined as units exhibiting a broader waveform (duration > the daily 10th percentile) and a lower firing rate (rate < the daily 90th percentile). Units that did not fall into these distinct quadrants were labeled as ‘unclassified’.

### Identification of putative inhibitory connections

Putative inhibitory monosynaptic connections, between pairs of co-recorded neurons, were identified by detecting significant, short-latency troughs in the firing-rate crosscorrelograms (CCGs) using a probabilistic thresholding approach, following previous procedures^29,71^. CCGs were computed with 0.67 ms bins over a ±66 ms window. A baseline CCG was computed by convolving raw CCGs with a hollowed gaussian kernel (σ = 16 ms). Only connections where the CCG contained at least 1,000 counts were further considered. The baseline CCG was subtracted from the raw CCG. Trough detection was then performed on the baseline-corrected CCG. Detection of inhibitory connections relied on a probabilistic thresholding approach. For each bin in the CCG, a lower confidence bound was calculated as the 1st percentile of the Poisson distribution derived from the predicted baseline. A neuron pair was classified as having a putative inhibitory connection only if the observed CCG trough fell below this 1% threshold. To ensure the detected suppression was robust to noise, the significant reduction in firing probability was required to persist for at least three consecutive bins (2 ms) within the 0.01–8 ms post-synaptic interval, thereby avoiding overlap with the zero-lag bin, which would suggest modulation by common input. Connection probability was obtained by dividing the number of identified connected pairs by the count of all pairs tested for a connection, for each developmental day independently.

### Rate maps and angular tuning curves

2D rate maps were generated by discretizing the environment into a grid of 33×33 bins. For each bin, the firing rate of each cell was computed as the number of spikes in the bin, divided by time spent in the bin. Rate maps were smoothed with a cross-validated smoothing procedure as described previously ^11^, using a kernel with a standard deviation of 1.75 bins. Unvisited bins (NaNs) within the map were filled using an inpainting algorithm.

Angular tuning curves with respect to head direction were calculated by binning the angular variable into 60 evenly spaced angular bins. For each 6° bin, the spike rate was calculated as the number of spikes divided by time spent in the bin. Angular tuning curves were smoothed with the same cross-validated smoothing procedure as the spatial rate maps (0.01 rad < *σ* < 1 rad).

### Desynchronization analysis

To quantify the coupling of activity in individual neurons to the global network state, we computed a population synchrony measure based on the correlation between individual unit activity and the total population firing rate.

Spike trains for all isolated units were first discretized into non-overlapping time bins. We assessed this across different temporal scales: 50, 100, 200 and 500ms.

For a given bin size, the population activity vector was constructed by summing the spike counts of all recorded units within each time bin. Specifically, if *ni(t)* represents the spike count of unit *i* in time bin *t*, the population vector *P(t)* was defined as:

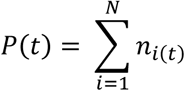

where N is the total number of units in the session.

To estimate the synchrony of a single unit to the network, we calculated the Pearson correlation coefficient (*r*) between that unit’s binned spike vector, *ni*, and the population activity vector, *P*. A session-wide score was then reported as the median of these correlation coefficients across all units in the recorded population.

The Spike Time Tiling Coefficient (STTC) was calculated as previously reported^43,72^, adapting routines from https://github.com/OpatzLab/HanganuOpatzToolbox. Briefly, correlations between spike trains were calculated as follows:

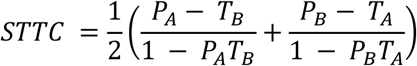

*PA* is he fraction of spikes in train A for which a corresponding spike from train B is present within the time interval ±Δ*t .* The proportion of the total recorded observation time that is covered (or ‘tiled’) by the ±Δ*t* temporal neighborhoods surrounding the spikes in train A. The variable ±Δ*t* functions as the temporal resolution parameter, setting the specific time window used for calculating the STTC, with the following time lags utilized: 50, 100, 200 and 500 ms.

### Histology and recording locations

The rat pups received a lethal dose of pentobarbital, after which they were perfused intracardially with saline followed by 4% formaldehyde. The brains were extracted and stored in 4% formaldehyde, and were later cut in 30–µm sagittal sections using a cryostat. The sections were Nissl-stained with cresyl violet. The probe shank traces were identified in photomicrographs. A map of the probe shank was aligned to the histology by using two reference points that had known locations in both reference frames: (1) the tip of the probe shank; and (2) the intersection of the shank with the brain surface. In all cases, the shank traces were near-parallel to the cutting plane, therefore it was deemed sufficient to perform a flat 2D alignment in a single section where most of the shank trace was visible. The aligned shank map was then used to calculate the anatomical locations of individual shanks.

Note that estimates of anatomical locations are subject to some measurement error owing to the limited accuracy of the alignment process and the fact that units may be detected some distance away from the recording site. In two animals (#29404 and #29056), the reconstruction of the final implant position (MEC or PaS) was not possible due to damage to the stained sections containing the probe shank traces.

### Data analysis and statistics

Data analyses were performed with custom-written scripts in Matlab and Python. Statistical analysis was done using Matlab and Python. Results are reported as median and interquartile range (IQR) unless otherwise indicated. Statistical comparisons between groups were performed using non-parametric methods to avoid assumptions of normality, unless otherwise indicated. For datasets with limited sample sizes (N < 5 per group) we calculated p values using exact permutation tests. For larger sample sizes, we employed the Mann–Whitney U test. The study did not involve any experimental subject groups, so random allocation and experimenter blinding did not apply and were not performed.

## Data availability

The datasets generated during the current study will be available at the time of publication. Source data are provided with this paper.

## Code availability

Code for reproducing the analyses in this article will be available at the time of publication.

## Author contributions

All authors planned and designed experiments, conceptualized and planned analyses; C.M.L. trained M.G. in implantation and recording procedures; M.G. and J.C. implanted animals and performed experiments; M.G., E.H. and J.C. wrote software and analyzed and visualized data, with input from all authors; M.G. curated data; M.G. and E.I.M. wrote the paper, with periodic inputs from M.B.M. and E.H.. All authors participated in discussion and interpretation. B.A.D., M.-B.M. and E.I.M. supervised the project. M.G., M.-B.M. and E.I.M. obtained funding for the project.

## Acknowledgements

We thank; We thank R.J. Gardner, D. Hayden, S. Ball, K. Haugen, E.H. Holmberg, K.J. Jenssen, E. Kråkvik, D.J. Hayden, and H. Waade for technical assistance; the veterinary staff for animal care; M.P. Witter for advice on implantation coordinates and assessment of recording locations; A.Z. Vollan, R.J. Gardner, R. Saxena, V. Olsen for discussions.

The work was supported by EMBO Postdoctoral Fellowships to M.G. (ALTF 738-2022), a Synergy grant to E.I.M. from the European Research Council (KILONEURONS, grant 951319); FRIPRO grants to E.I.M. (grant 286225) and M.-B.M. (grant 300394/H10) from the Research Council of Norway; two Centre of Excellence grants to M.-B.M. and E.I.M. (Centre of Neural Computation, grant 223262, and Centre for Algorithms in the Cortex, grant 332640) from the Research Council of Norway; a National Infrastructure grant to E.I.M. and M.-B.M. from the Research Council of Norway (NORBRAIN, grant 295721 and 350201); a grant from the Kavli Foundation to M.-B.M. and E.I.M.; a grant from the Research Council of Norway (iMOD, NFR grant #325114) to B.A.D.; a gift from E. Solberg and family; and a direct contribution to M.-B.M. and E.I.M. from the Ministry of Education and Research of Norway.

---

**Supplementary Table 1.** Information for each animal. Top: semichronic animals P8–P12; middle: chronic animals P8–P12; bottom: chronic animals P15–P19. In semi-chronic animals the surgery, recording, and perfusion were completed within a single day. In chronic animals the recordings extended over multiple days, and pups were returned to the nest at the end of each recording.

| Animal code | Age (at implant) | Weight (at implant) | Sex | # Units (per day) | Toroidal Ensembles | Shanks | Implant position |
| --- | --- | --- | --- | --- | --- | --- | --- |
| 29387 | P8 | 21g | Male | 427 | 0 | 2,3 | L2/L3, PaS |
| 29404 | P8 | 21g | Male | 619 | 0 | 1, 2 | n/a |
| 29501 | P8 | 20g | Male | 350 | 0 | 1 | PaS |
| 29508 | P8 | 21g | Male | 753 | 0 | 1 | L2/L3, PaS |
| 29416 | P9 | 20g | Male | 1102 | 0 | 1, 2 | PaS |
| 29468 | P9 | 21g | Male | 572 | 0 | 2 | PaS |
| 29541 | P9 | 20g | Male | 353 | 0 | 2, 3 | L2/3 |
| 29729 | P9 | 21g | Male | 469 | 0 | 1, 2 | L2/3, PaS |
| 28854 | P10 | 23g | Female | 905 | 0 | 3,4 | L2/3, PaS |
| 29298 | P10 | 23g | Male | 983 | 0 | 2,3 | L2/3, PaS |
| 28801 | P10 | 28g | Male | 1099 | 1 | 2,3,4 | L2/3, PaS |
| 28793 | P11 | 35g | Male | 1244 | 2 | 1,2,3,4 | L2/3 |
| 28901 | P11 | 29g | Male | 1182 | 1 | 2,3,4 | L2/3 |
| 29301 | P11 | 35g | Male | 1181 | 1 | 1,2 | L2/3, PaS |
| 28594 | P12 | 38g | Male | 1234 | 0 | 1,2 | PaS |
| 29023 (Mirtillo) | P10-12 | 26g | Male | 1186; 948; 1341 | 2;1;2 | 2,3,4 | L2/3, PaS |
| 29078 (MT) | P10-12 | 26g | Female | 767; 646; 643 | 1;1;0 | 1 | L2/3 |
| 29316 (Taro) | P10-16 | 25g | Male | 1246; 1057; 1158 | 2;2;3 | 2,3 | L2/3 |
| 28738 (Amarena) | P14-19 | 36g | Male | 1240 - 701 | 1;1;0;1;2 | 4 | L2/3 |
| 28761 (Vanilla) | P14-19 | 37g | Female | 866 - 1145 | 2;1;1;1;1 | 3, 2/4 | L2/3, PaS |
| 29056 (Cookie) | P14-19 | 40g | Male | 954 - 1027 | 2;0,1;1;1 | 1,2,3,4 | n/a |
| 28697 (Caramello) | P18-19 | 51g | Male | 1289-1207 | 1;0 | 2,3,4 | L2/3 |
| 28723 (Cacao) | P16 | 44g | Male | 779 | 2 | 2,3,4 | L2/3, PaS |

## Extended Data

**Extended Data Fig. 1:**
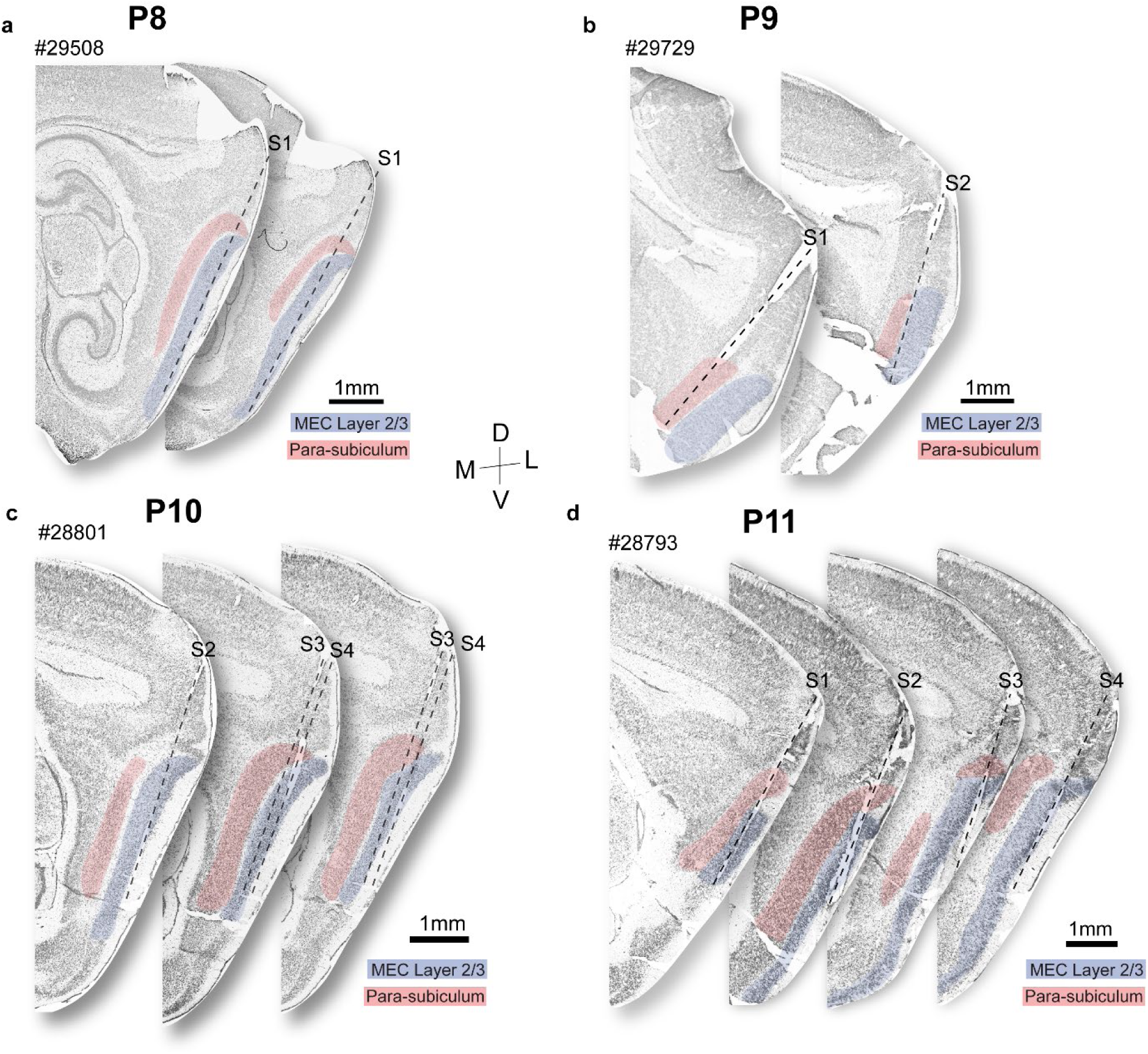
Representative histological localization of recording locations. Recording locations from representative animals across developmental ages. **a.** Serial sagittal sections of a representative animal at P8 (#29508). Sections are displayed from medial (left) to lateral (right). Sections with the clearest tracks for each shank selected for the recording (indicated by number) are shown. Regions are indicated with shaded colors: MEC Layer 2/3 in blue, PaS in red. Toroidal clusters were not found in this animal (Supplemetary Table 1). **b.** Same as in (a) but for a representative animals at P9 (#29729). Toroidal clusters were not found in this animal. **c.** Same as in a–b but for a representative animal at P10 (#28801). A toroidal cluster was found in this animal. **d.** Same as in a–c but for a representative animal at P11 (#28793). A toroidal cluster was found in this animal.

**Extended Data Fig. 2:**
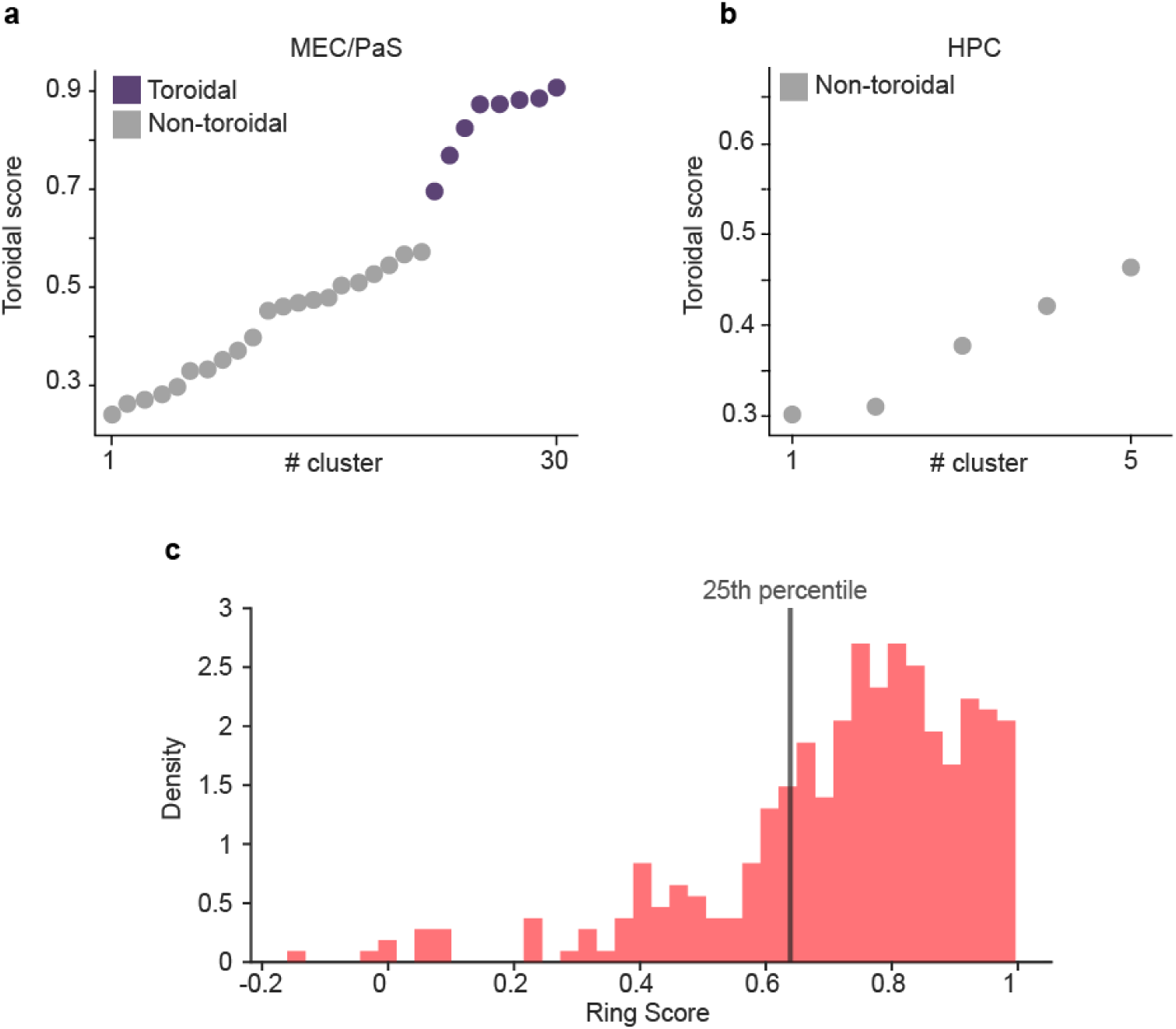
Quantification of toroidal scores and ring scores in adult animals. **a.** Toroidal scores for individual clusters in the adult MEC–PaS (n=5 animals), from a published dataset^29^. To reliably identify toroidal topology, a threshold of 0.60 was established based on the largest gap in the data distribution. Each dot represents a cluster; purple indicates clusters classified as toroidal, and grey indicates non-toroidal clusters. **b.** Toroidal scores for hippocampal (HPC) clusters (n=2 animals). No clusters exceeded the 0.60 threshold for toroidal classification. **c.** Distribution of ring scores for individual head direction cells in adult animals (n=5 animals; inclusion criteria: mutual information score between the animal’s head direction and the neuron’s firing activity > 1 bit). The solid vertical line marks the 25th percentile of the distribution, corresponding to a ring score of 0.63. This score was used as a threshold for identifying cells with tuning to the ring from P8–P12.

**Extended Data Fig. 3:**
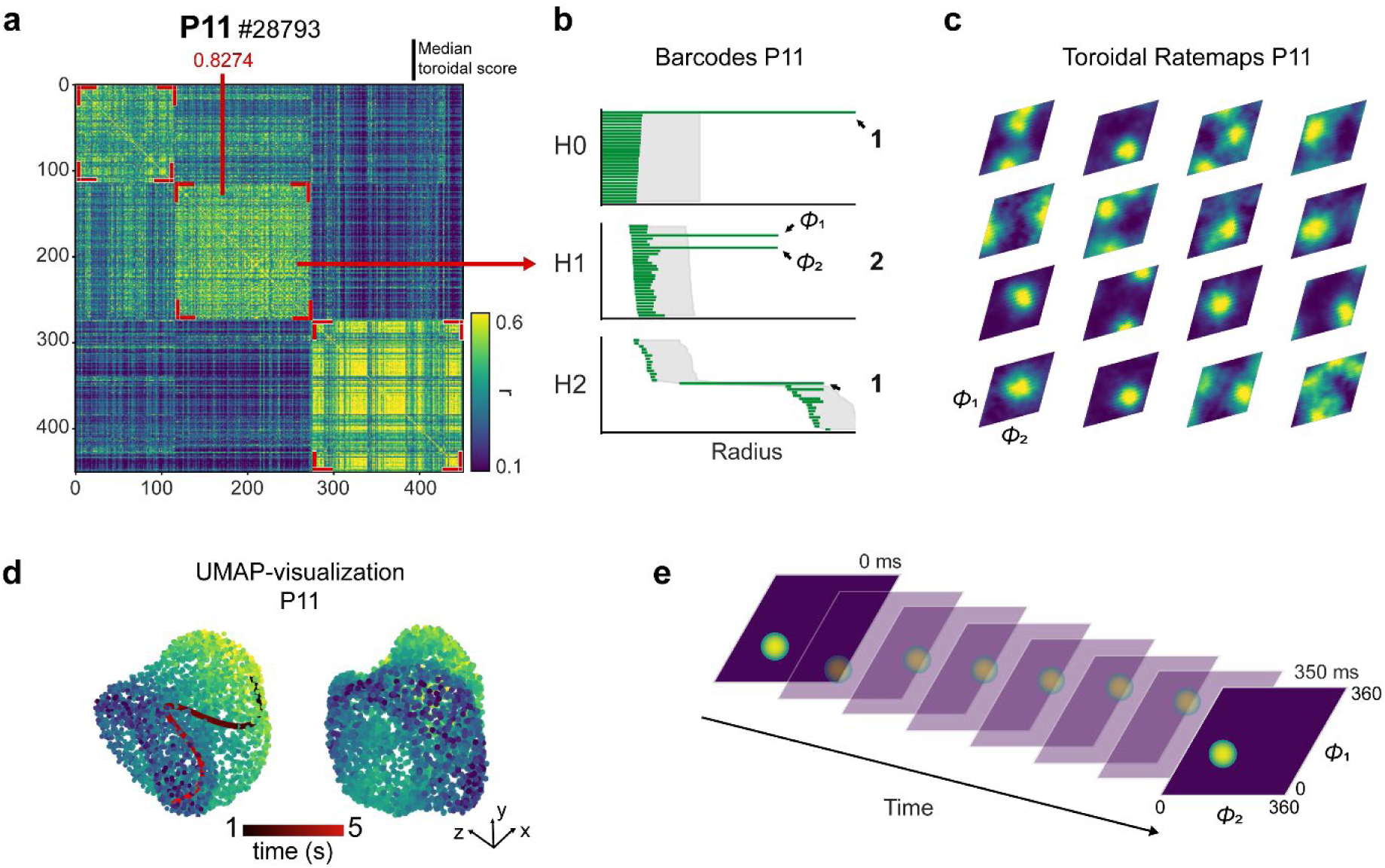
Toroidal activity at P11. **a.** Correlation distance matrix of the time-lagged cross-correlation of the firing rate activity of all recorded cells in a rat pup recorded at P11. Highlighted with red arrows is a cluster with high median toroidal score. **b.** Persistent cohomology analyses for the P11 ensemble highlighted in panel **a**. Barcodes indicate the homological signature of a 2-D torus: one 0-D bar (*H*^0^), two 1-D bars (*H*^1^) and one 2-D bar (*H*^2^). Grey shading indicates the longest lifetimes among 100 iterations in shuffled data (aligned to lower values of original bars; P<0.01). **c.** The majority of cells in the cluster highlighted in panel **a**. have toroidal firing fields, meaning that their activity is tuned to a restricted phase of the torus. The example shows 16 representative cells with tuning to decoded toroidal positions (*ϕ*1 and *ϕ*2 in H1). **d.** UMAP projection reveals a 3D toroidal point cloud in the population activity of the ensemble highlighted in panel **a.**, with colors representing loading on the first principal component (time bins were subsampled taking every 65th time bin, resulting in 650–ms intervals between selected samples; see Methods). Bold black/red line represents a 5-s trajectory in the point cloud to the left, showing smooth movement over the toroidal manifold. **e.** Representative example of toroidal decoding of the P11 data in a–d over time. Each circular parametrization was interpolated to the population activity to obtain the decoded toroidal coordinates (computed by finding the mass center of the summed distributions, weighted by the population activity vector to be decoded). Successive planes show the activity peak across consecutive time intervals (50ms time bins) over the two 1-D bars in the H1 dimension (*ϕ*1 and *ϕ*2).

**Extended Data Fig. 4:**
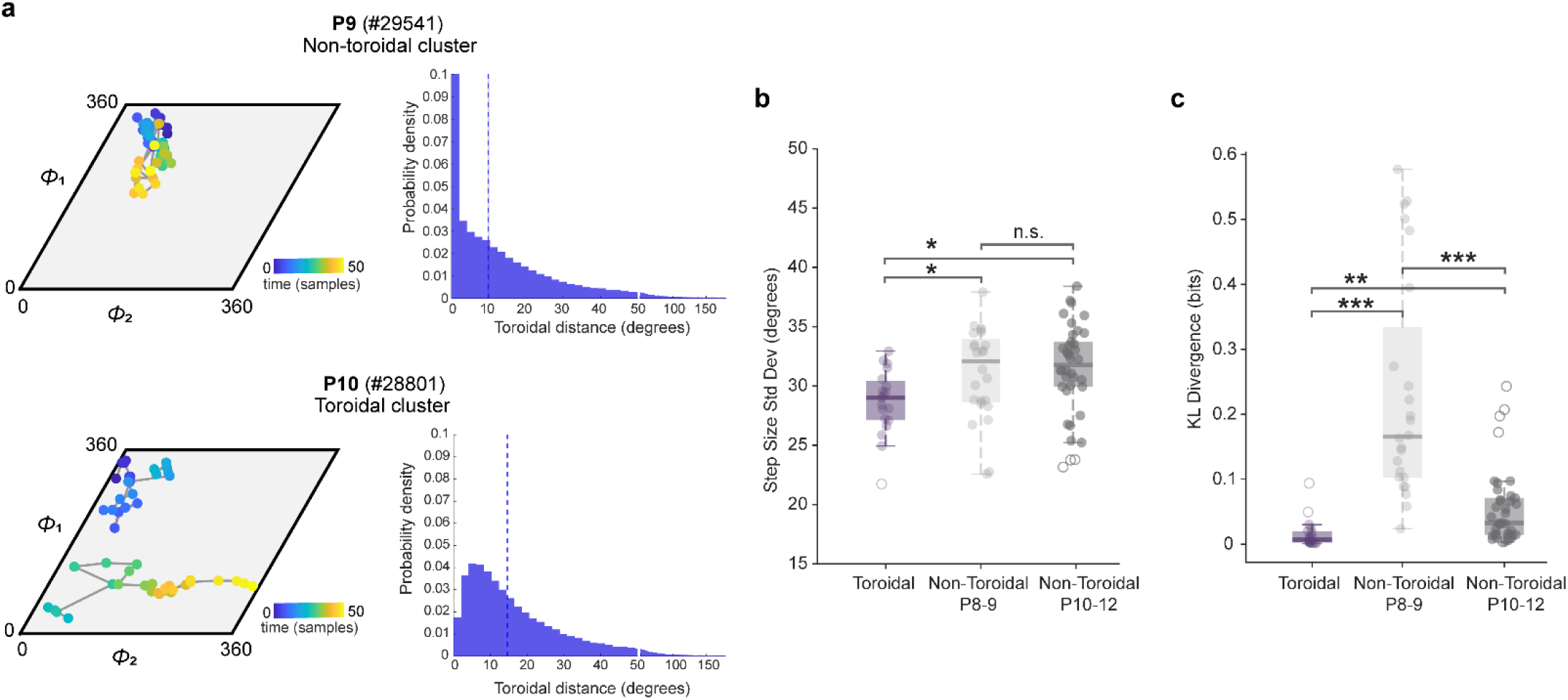
Toroidal clusters exhibit smoother movement dynamics than non-toroidal clusters. **a.** Representative movement dynamics for a non-toroidal cluster at P9 (top) and a toroidal cluster at P10 (bottom, using the same clusters as in Fig.1). Left panels: trajectories of the decoded bump position projected onto the toroidal manifold, displayed as a rhombus. The color gradient indicates time progression over a 2.5 s window (50 steps). Grey lines connect consecutive time points; breaks in the lines indicate crossings of the periodic boundaries. Right panels: Probability density functions of step sizes (phase shifts in degrees) between consecutive time points, computed as the shortest geodesic path on the hexagonal torus (minimum across 9 periodic replications), with periodic boundary conditions accounted for. At P9 step sizes are heavily concentrated near zero, indicating predominantly small, localized movements. Distributions are shown for time lags of 50ms. The x axis scale changes at 50 ms to highlight better the differences at small step sizes. Dashed vertical line indicates median step size. **b.** Quantification of trajectory smoothness expressed as the standard deviation of step sizes on the manifold (distances between consecutive decoded positions on the two longest H1 dimensions, regardless of whether they met the criteria for a torus). A Kruskal-Wallis test revealed significant differences in step size standard deviation between toroidal groups and non-toroidal groups at P8-9 and P10-12 (χ²(2) = 9.12, P = 0.01). Toroidal clusters at P10–12 (purple) exhibited significantly lower step-size variability than both non-toroidal clusters from younger animals (P8–9, light grey; Bonferroni correction, p = 0.04) and age-matched non-toroidal clusters (P10–12, dark grey; Bonferroni correction, p = 0.01). The two non-toroidal groups did not differ significantly (Bonferroni correction, p = 1.00). Central mark indicates median; box edges represent interquartile range. The length of the whiskers indicates 1.5 times the interquartile range. Outliers are represented by empty circles. n.s., non-significant; *, P<=0.05. The results indicate that activity on toroidal manifolds encodes continuous, smooth trajectories, whereas activity decoded from non-toroidal clusters produces more erratic, discontinuous jumps. **c.** Kullback-Leibler (KL) divergence (in bits) for distributions of step-size in toroidal and non-toroidal clusters (P8-9 and P10-12). KL divergence is significantly different between the three sets of clusters (Kruskal-Wallis test, χ²(2) = 44.63, p = 10^−10^). Non-toroidal clusters at P8–9 (Bonferroni correction, p = 10^−10^) and P10–12 (Bonferroni correction, p = 0.009) exhibit significantly higher divergence compared to the P10–12 toroidal group. Divergence is also significantly higher in non-toroidal clusters at P8–9 compared to non-toroidal clusters at P10–12 (Bonferroni correction, p = 10^−5^), indicating distinct movement statistics across the three groups. Central mark indicates median; box edges represent interquartile range. The length of the whiskers indicates 1.5 times the interquartile range. Outliers are represented by empty circles; **, P<=0.01 ***, P<=0.001.

**Extended Data Fig. 5:**
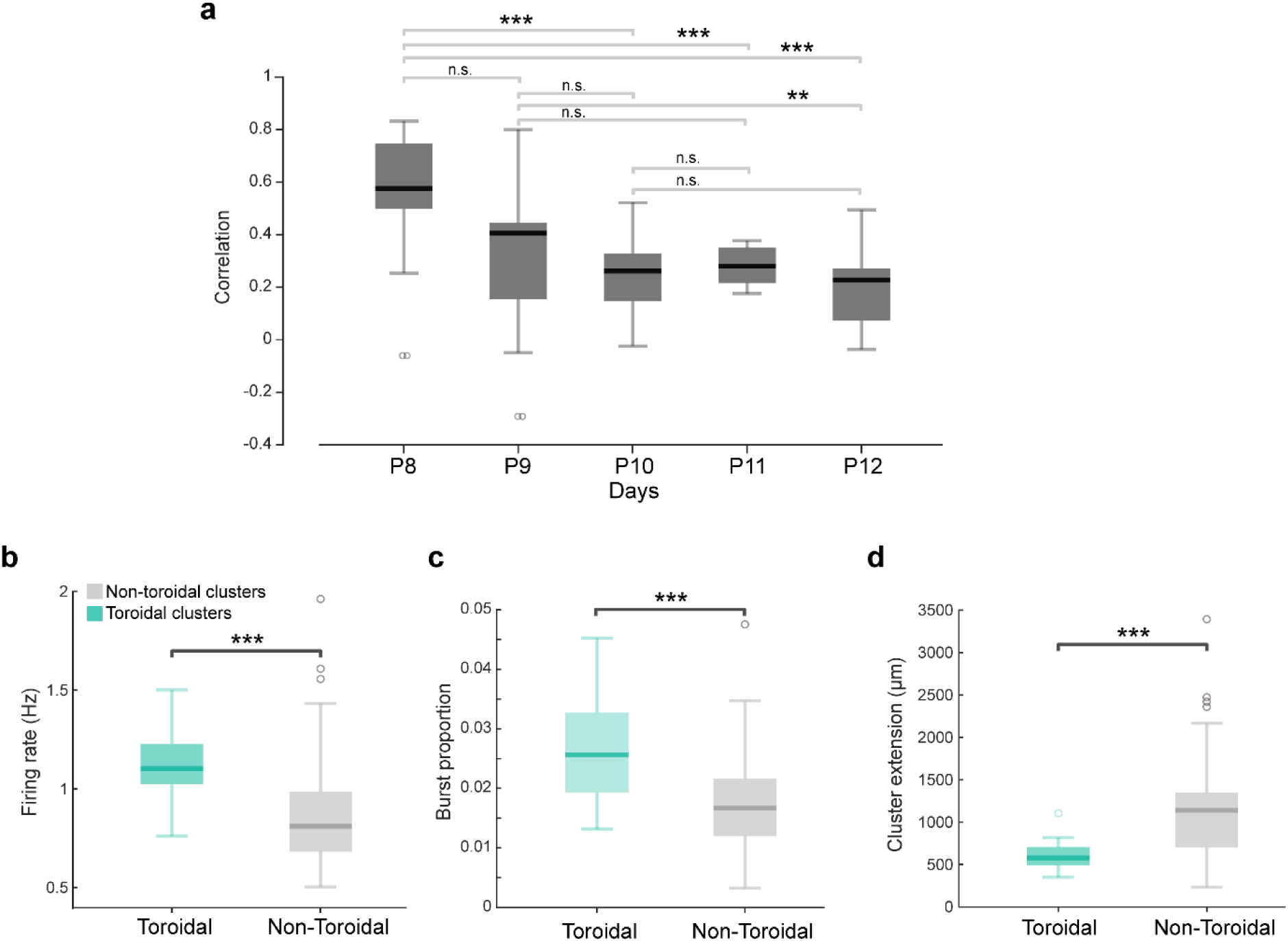
Characteristics of toroidal clusters across postnatal days. **a.** Average correlation scores comparing all simultaneously recorded cell pairs (both within and between clusters) of time-lagged cross-correlation matrices (as in Fig. 1c) from P8 to P12. Pairwise correlation scores decrease as postnatal age increases, resulting in a negative correlation between the two variables (Spearman correlation r = -0.357, P = 10⁻⁸). A Kruskal-Wallis test revealed significant differences among groups (χ²(4) = 42.96, P = 10⁻⁸). Pairwise post-hoc comparisons were performed using a Bonferroni correction and significant comparisons are reported in the plot. Central lines indicate the median; box edges represent the interquartile range (IQR). Whiskers extend to a maximum of 1.5 times the IQR. Outliers are indicated by empty circles. n.s., non-significant; **, P ≤ 0.01; ***, P ≤ 0.001. **b.** Firing rate (Hz) for toroidal and non-toroidal clusters (P10–P12 and P8–P12, respectively). Toroidal clusters display higher firing rates [1.10 (1.02–1.22), median and IQR] than non-toroidal ones [0.81 (0.6823–0.9865), median and IQR; Mann-Whitney U test, Z = 3.95; p value =0.0001]. Central mark indicates median; box edges represent interquartile range. The length of the whiskers indicates 1.5 times the interquartile range. Outliers are indicated by empty circles. ***, P<=0.001. **c.** Burst proportion in toroidal versus non-toroidal clusters (P10–P12 and P8–P12, respectively). Burst proportion is computed as the number of discrete burst events (defined as groups of spikes with inter-spike intervals under 7 ms) divided by the total number of firing events encompassing both bursts and single spikes. Toroidal clusters show a higher proportion of burst firing [0.025 (0.01–0.03), median and IQR] than non-toroidal clusters [0.016 (0.01–0.02), median and IQR) ; Mann-Whitney U test, Z = 3.82, p value = 0.0001]. Central mark indicates median; box edges represent interquartile range. The length of the whiskers indicates 1.5 times the interquartile range. Outliers are represented by empty circles. ***, P<=0.001. **d.** Anatomical extension (measured in µm) along the silicon probe shanks for toroidal clusters (green) and non-toroidal clusters (grey). The extension of a cluster is estimated by extracting the estimated depth from all neurons within the cluster and then computing the difference between the 95th percentile and the 5th percentile of these depth values (thus mitigating the influence of extreme spatial outliers). Toroidal clusters are confined within a shorter region [577.49 (492.51–705.46), median and IQR] compared to non-toroidal clusters [1139.95 (702.34–1347.53), median and IQR; Mann-Whitney U test, Z = -3.91, p value =0.0001]. Central mark indicates median; box edges represent interquartile range. The length of the whiskers indicates 1.5 times the interquartile range. Outliers are represented by empty circles; ***, P<=0.001.

**Extended Data Fig. 6:**
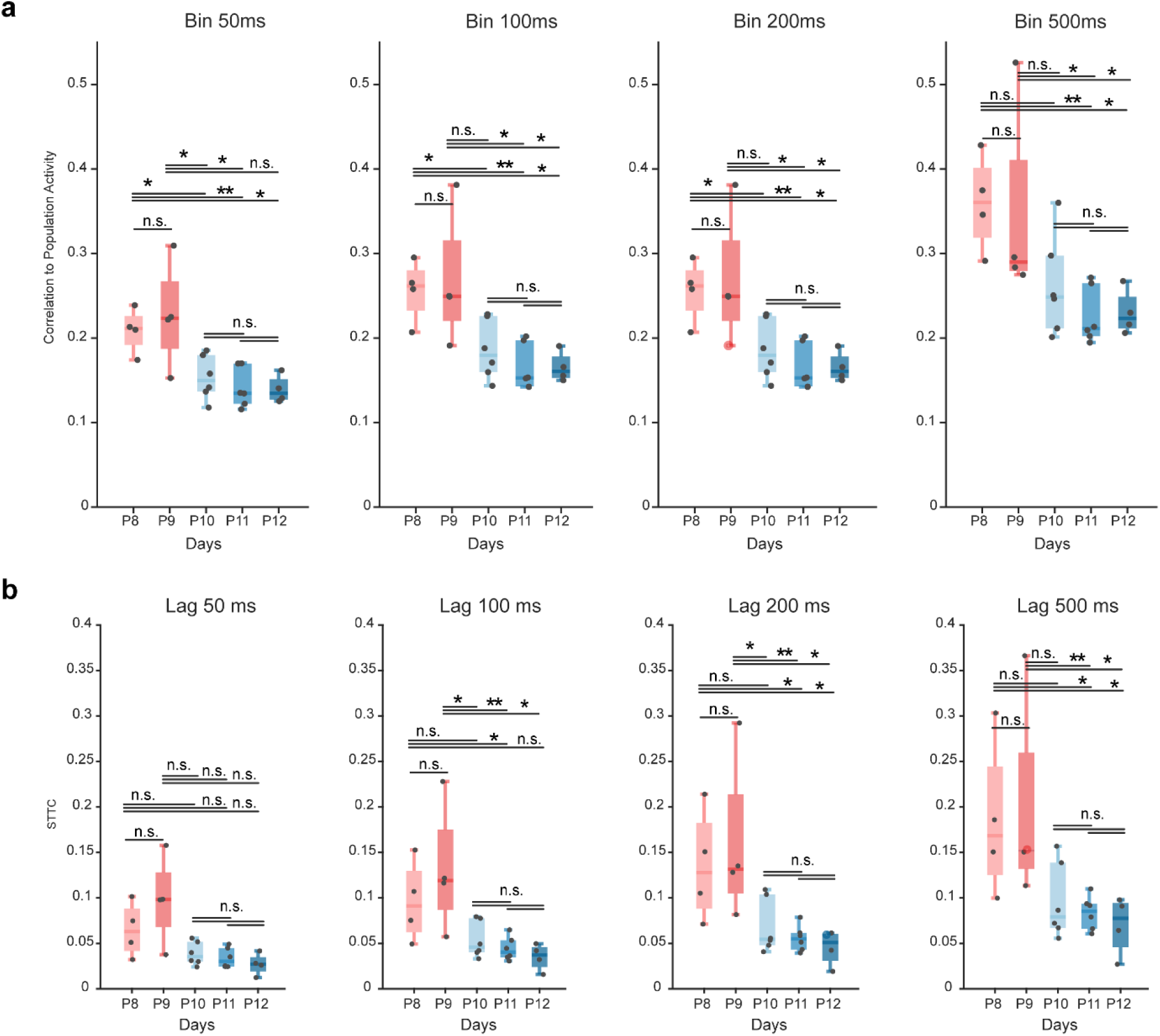
MEC network dynamics across temporal binning parameters. **a.** Correlation between spike activity of individual neurons and population activity across developmental ages and time bins (time bins: 50ms, 100ms, 200ms and 500ms). Data pooled across 18 animals (n = 24 sessions). Spearman correlation (from left to right): r values = -0.6891, -0.7184, -0.7095, -0.6908; p = 0.0002, 0.0001, 0.0001, 0.0002. Comparisons across days were performed using an exact permutation test, followed by a Benjamini-Hochberg false discovery rate (FDR) correction for multiple comparisons. Central mark indicates median; box edges represent interquartile range. The length of the whiskers indicates 1.5 times the interquartile range. n.s., non-significant; *, P<=0.05; **, P<=0.01, significant after FDR correction. **b.** Spike Time Tiling Coefficient (STTC) scores from P8 to P12. The STTC assesses pairwise correlation by calculating the proportion of spikes in one train that fall within a specific time lag of spikes in another. Unlike standard correlation measures, the STTC normalizes these overlaps against the total time ‘tiled’ by the windows, making the metric robust to fluctuations in firing rate (see Methods). Time lags: 50ms, 100ms, 200ms and 500ms. Data pooled across 18 animals (n = 24 sessions. Lower STTC values at P10-12 indicate higher decorrelation in the neural population. Spearman correlation (from left to right): r values = -0.6162, -0.6802, -0.6917, -0.6757; p = 0.0007, 0.0001, 0.0001, 0.0001). Comparisons across days were performed using an exact permutation test, followed by a Benjamini-Hochberg false discovery rate (FDR) correction for multiple comparisons. Central mark indicates median; box edges represent interquartile range. The length of the whiskers indicates 1.5 times the interquartile range. n.s., non-significant; *, P<=0.05; **, P<=0.01, significant after FDR correction.

**Extended Data Fig. 7:**
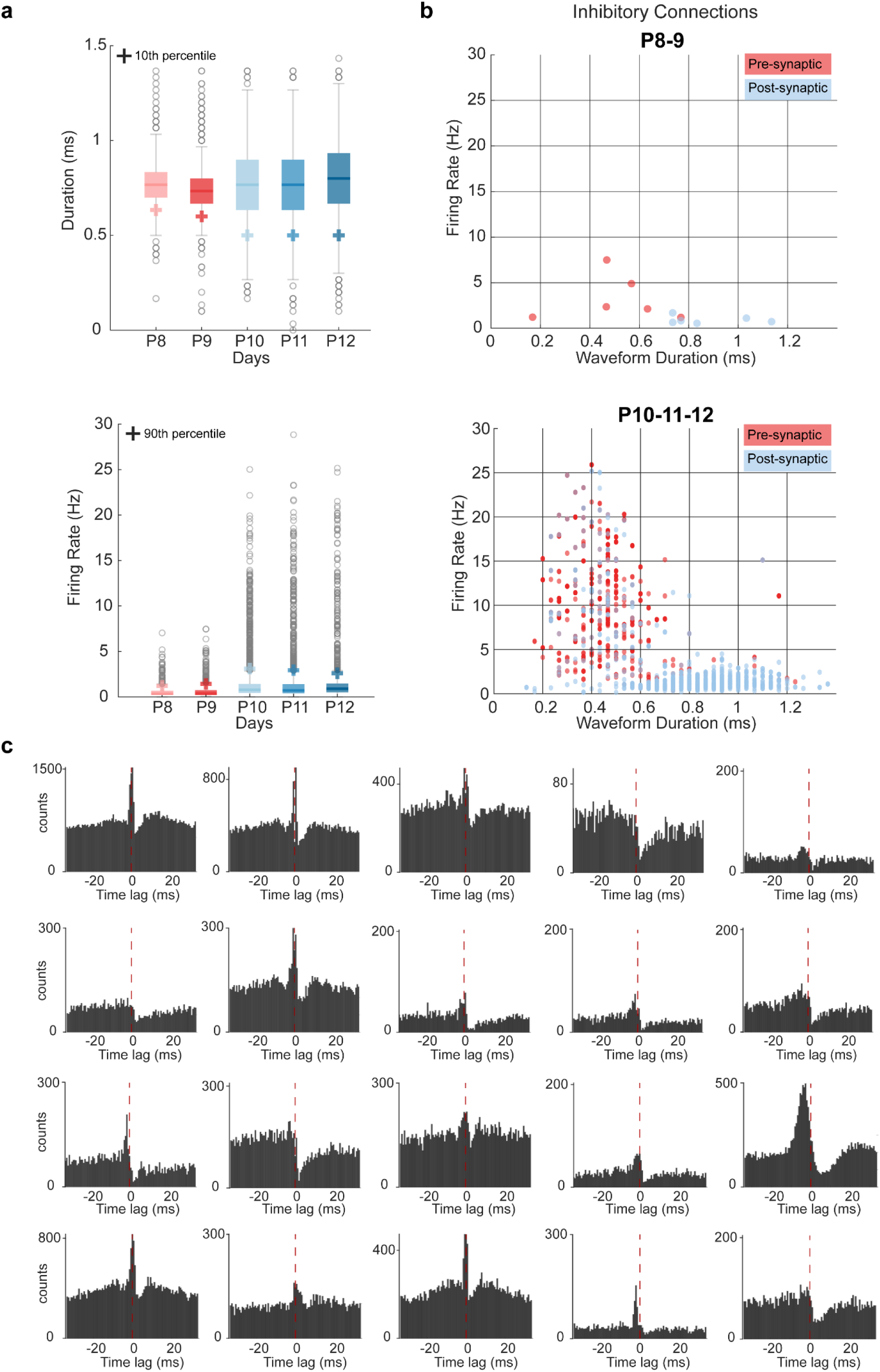
Emergence of fast-spiking cells and putative inhibitory connections. **a.** Development of waveform duration and firing rates for all individual units in MEC–PaS between P8 and P12. Top panels: distributions of average waveform width become progressively narrower (10^th^ percentiles decrease from 0.63ms at P8 and 0.60ms at P9 to 0.50ms at P10–P12). Bottom panel: firing rates become progressively higher across days (95^th^ percentile values increase from 1.2 Hz at P8 and 1.4 Hz at P9 to 3.10 Hz at P10, 2.9 Hz at P11 and 2.5 Hz at P12). Central mark indicates median; box edges represent interquartile range. The length of the whiskers indicates 1.5 times the interquartile range. Outliers are indicated by empty circles. **b.** Among all detected cell pairs, the majority of pre-synaptic cells have a short-latency waveform and high firing rate. Scatter plot compares average firing rate (y-axis) and waveform half-width duration (x-axis) for presynaptic (red) or postsynaptic (light blue) cells of a cell pair at P8–9 (top) and P10–12 (bottom). Note the substantial increase in number of identified connections between the two age groups. **c.** Additional examples of putative inhibitory connections at P10–P11. The sharp trough near zero latency indicates a suppressed firing probability of the post-synaptic neuron immediately following a pre-synaptic spike, consistent with an inhibitory connection between the cell pairs.

**Extended Data Fig. 8:**
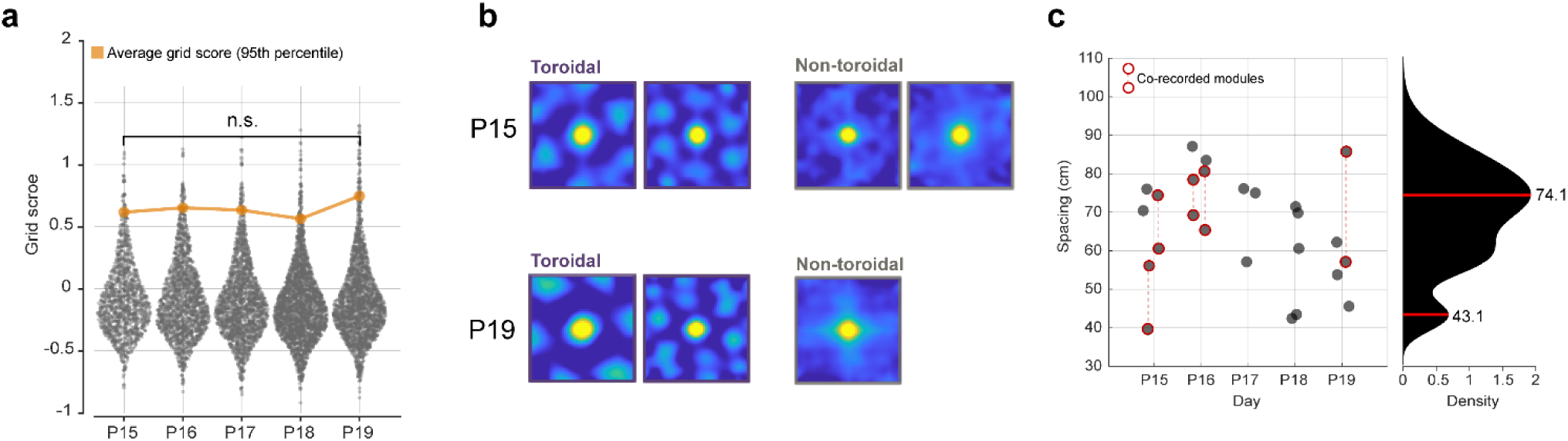
Grid patterns in toroidal and non-toroidal clusters. **a.** Distribution of grid scores across all cells in non-toroidal clusters. Each dot represents one unit (from 1,336 to 2,792 per day). Grid scores did not increase but decreased between P15 and P18 (Kruskal-Wallis test, χ^2^= 45.66, p< 10^−7^). No difference was observed between the scores on the first (P15) versus the last day (P19) of exploration (Dunn’s Post-Hoc test: P15 vs P19: p-value = 0.545, diff = 180.21). Orange line shows 95^th^ percentile values (mean across animals). **b.** Representative spatial autocorrelograms for neurons within toroidal and non-toroidal clusters, simultaneously recorded from the same animal (#28761) at P15 and P19. Two toroidal clusters (left panels) exhibit clear hexagonal grid-like periodicity. Note that these clusters differ in grid spacing, consistent with the early emergence of discrete scales. **c.** Distribution of grid spacing for all detected toroidal clusters from P15 to P19 (left panel). Each dot represents mean spacing of one module. Red circles connected by dashed vertical lines indicate simultaneously recorded modules within the same session. The kernel smoothed density (KSD) estimate of the distribution (right panel) reveals a bimodal distribution of spacing across the population, with distinct peaks centered at approximately 43.1 cm and 74.1 cm.

**Extended Data Fig. 9:**
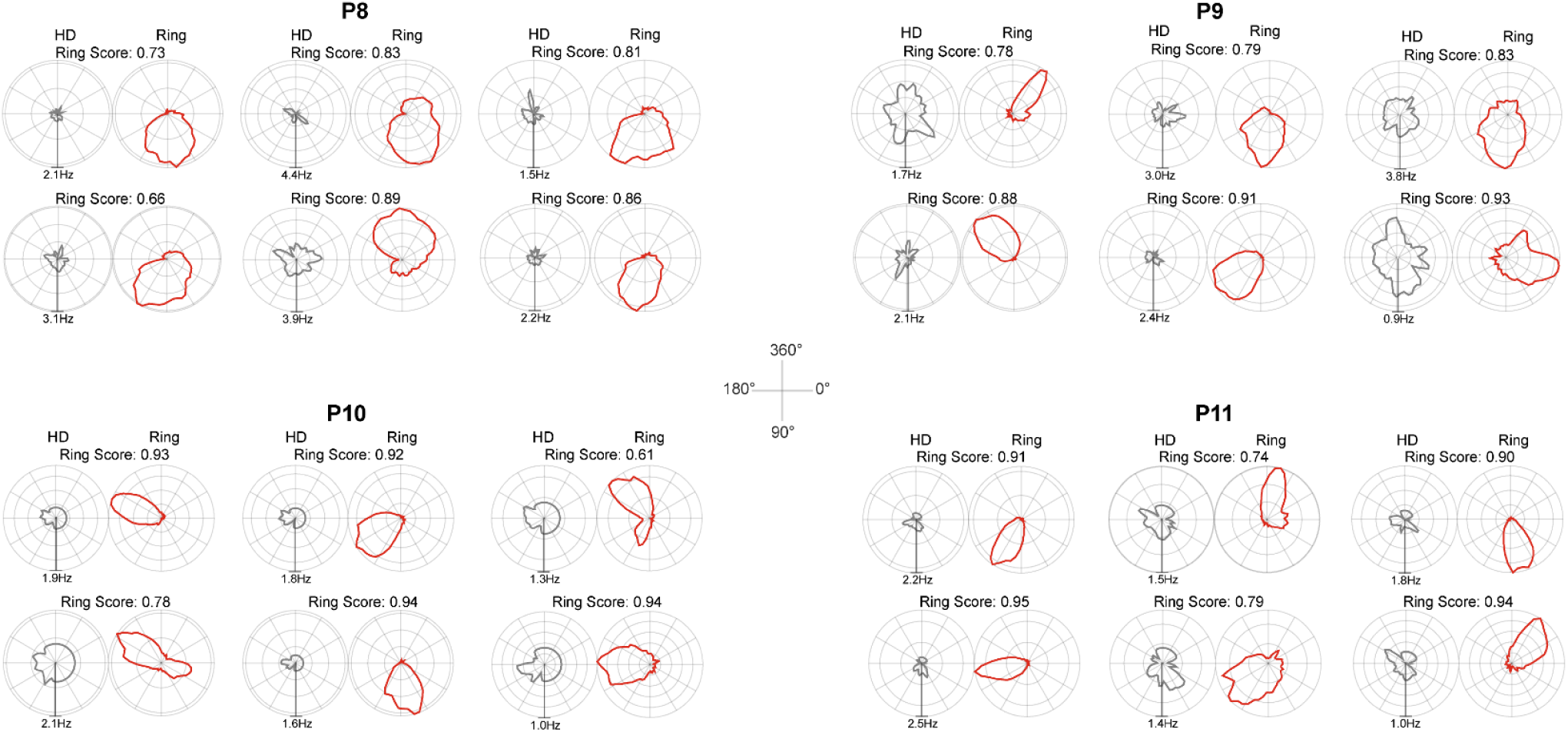
Representative examples across ages of individual cells tuned to an intrinsic ring-like manifold. Representative polar tuning profiles of single units recorded from P8 to P11 (left to right, different animals). Each pair compares the unit’s tuning to the external head direction (left, grey) versus its tuning to inferred position on the internally organized ring manifold (right, red). Ring coordinates for decoding were inferred for all clusters from the longest bar in the H1 dimension. The radial axis represents the firing rate (Hz). Plots are peak-scaled for each unit. The angular axis represents direction (0°–360°) plotted clockwise. Ring scores are reported for each cell. Note that sharp tuning to the internal ring manifold is present as early as P8.

**Extended Data Fig. 10:**
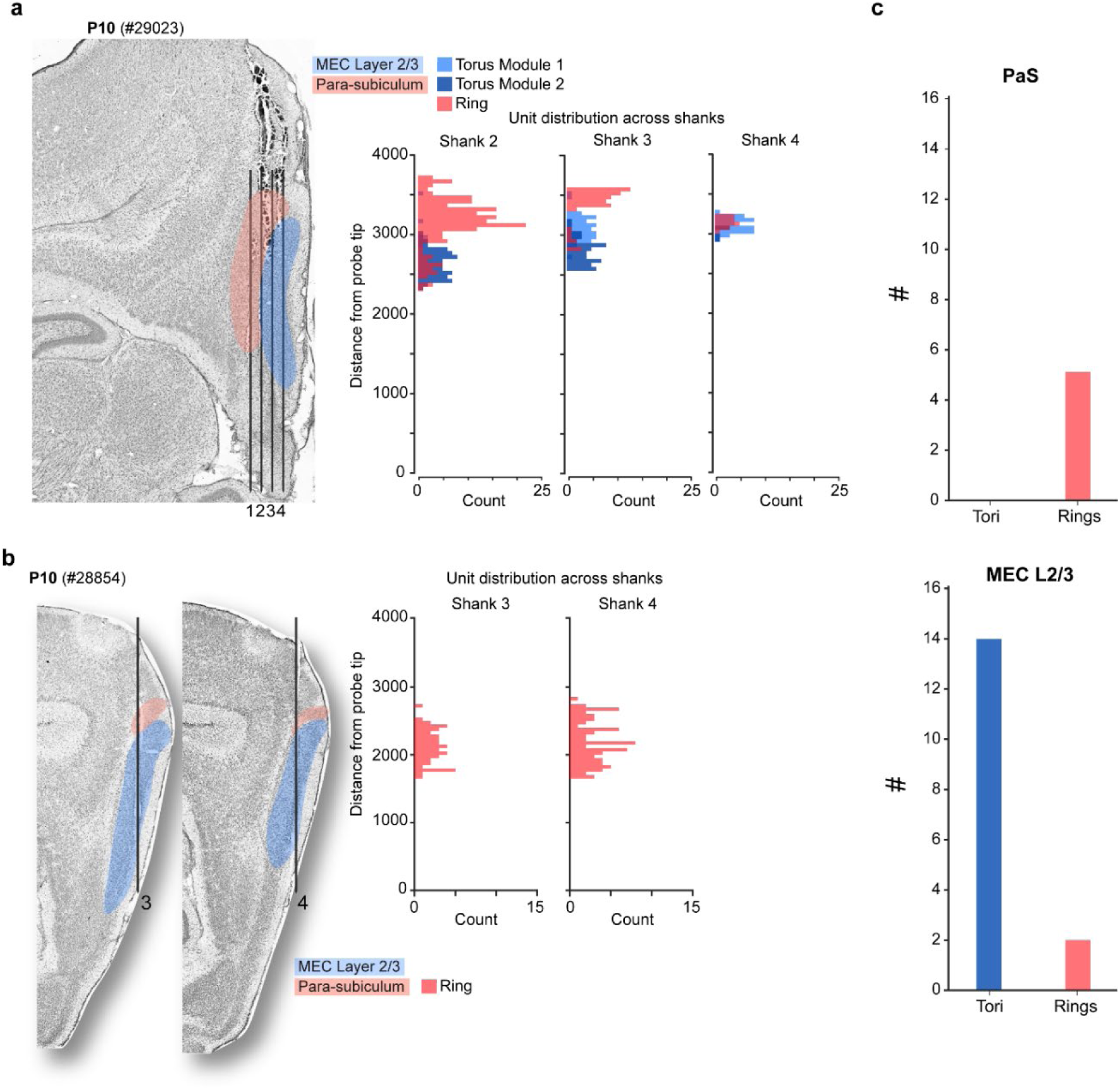
Ring and torus manifolds are located in PaS and MEC L2/3, respectively. **a.** Histological localization of cells belonging to individual clusters in the time-lagged cross-correlation analysis. Left: Nissl-stained sagittal section showing the tracks of a four-shank Neuropixels probe spanning the PaS (highlighted in light red) and MEC L2/3 (highlighted in light blue) on P10. Right: histograms showing distribution of recorded units along individual probe shanks (y-axis: distance from ventral probe tip), color-coded by inferred cluster topology (red: ring; light and dark blue: torus modules M1 and M2, respectively). **b.** Same as in **a.** but for an animal with a ring cluster detected in MEC on P10. **c.** Quantification of cluster topology by anatomical region. Bar plots display the total count of identified toroidal (blue) and ring (red) manifolds located within the histologically reconstructed PaS (left) and MEC L2/3 (right) regions. Ring manifolds were mostly observed in the PaS (5 out of 7 clusters), while toroidal manifolds were solely restricted to MEC L2/3. The two MEC rings were observed at P10 in animals that did not yet have toroidal clusters, suggesting that ring manifolds may appear temporarily as precursors to toroidal manifolds.

